# Antimicrobial resistance gene prevalence in a population of patients with advanced dementia is related to specific pathobionts

**DOI:** 10.1101/783969

**Authors:** Aislinn D. Rowan-Nash, Rafael Araos, Erika M.C. D’Agata, Peter Belenky

## Abstract

**Background:** The issue of antimicrobial resistance continues to grow worldwide, and long-term care facilities are significant reservoirs of antimicrobial-resistant organisms, in part due to high frequency of antimicrobial use. Patients with advanced dementia are particularly vulnerable to multidrug-resistant organism acquisition and antimicrobial overuse, which has negative consequences for the gut microbiome and can contribute to the selection and propagation of antimicrobial resistance genes. In this study, we longitudinally examined a group of advanced dementia patients treated with the fluoroquinolone antimicrobial levofloxacin, finding a correlation between abundance of pathogens and antimicrobial resistance genes, which we confirmed in a larger cohort of subjects with advanced dementia.

**Results:** We observed significant inter- and intra-subject heterogeneity in the composition of the microbiota of the longitudinal levofloxacin cohort, suggesting temporal instability. Within this dataset, we did not find significant impacts of levofloxacin on the diversity, composition, function, or resistome of the gut microbiota of this population. However, we were able to link the antimicrobial resistance gene burden in a sample with the relative abundance of several pathobionts – particularly *Escherichia coli*, *Proteus mirabilis*, and *Enterococcus faecalis*, as well as less-prevalent species including *Providencia stuartii* and *Staphylococcus haemolyticus*. Furthermore, we used metagenomic assembly and binning to demonstrate that these species had higher genomic resistance gene levels than common gut commensals, and we were able to predict antimicrobial resistance gene burden from the relative abundances of these species in a separate, larger cohort from the same population.

**Conclusions:** We found that the relative abundances of several pathobionts were correlated with and were even predictive of the level of antimicrobial resistance genes in corresponding samples, and that these species carried high levels of resistances genes in their assembled genomes. In order to test this observation, we utilized a larger metagenomics dataset from a similar population and confirmed the association between pathobiont abundance and antimicrobial resistance genes. Given the high frequency with which these species were found at high levels in this population and the underlying vulnerability to infection with multidrug resistant organisms of advanced dementia patients, attention to microbial blooms of these species may be warranted. Additionally, in this study, we were able to utilize genomic assembly from metagenomic data to more definitively associate antimicrobial resistance gene levels with specific assembled species; as this technology continues to develop, assembly could prove to be a valuable method to monitor both specific resistance genes and blooms of multidrug-resistant organisms.

## BACKGROUND

It is well-recognized that there is a growing threat of antimicrobial-resistant (AMR) bacterial strains that threaten the health and lives of millions worldwide. In the United States alone, the Centers for Disease Control and Prevention estimates that at least 2 million people get an AMR infection each year, and at least 23,000 die as a result[1]. A number of factors have driven the rise in AMR bacteria worldwide, including overprescription of antibiotics in the healthcare setting, over-the-counter access to antibiotics in some countries, and widespread use of antibiotics in animal husbandry for non-veterinary purposes[2–4]. Concerningly, hospitals and other medical institutions are frequent sites of AMR bacteria acquisition, where patients may already be ill or immunocompromised, antimicrobial use is common, and patient-to-patient transmission of AMR isolates can occur via inadequate hygiene or environmental contamination[5–8]. For example, AMR bacteria are highly prevalent in nursing homes, with estimates that over 35% of nursing home residents are colonized with multidrug resistant organisms (MDROs)[9–14]. This is particularly problematic in light of the fact that elderly patients in long-term care facilities may be frequently hospitalized, potentially serving as a mode of bidirectional transport of MDROs between healthcare facilities[15–17]. They are also prone to infections and are frequently treated with antimicrobials[18–20], which has long been associated with acquisition of MDROs and may not always be indicated[12, 17, 20–28].

The problem of MDRO prevalence and inappropriate antimicrobial use is of particular relevance in elderly subjects with advanced dementia, a population which receives extensive antimicrobial treatment which becomes more frequent closer to death, calling its benefit and effectiveness into question[28, 29]. Accordingly, advanced dementia specifically has been shown to be a risk factor of MDRO colonization[13, 30]. To examine this issue, the Study of Pathogen Resistance and Exposure to Antimicrobials in Dementia (SPREAD) was undertaken from 2009-2012 in order to analyze MDRO acquisition and appropriateness of antimicrobial prescription in elderly adults with advanced dementia residing in nursing homes[31]. Supporting the widespread nature of MDRO carriage in this population, analysis of SPREAD subjects revealed that there were significant baseline levels and new acquisitions of MDROs, and that there was notable spread of MDRO strains within and even between nursing home facilities[28, 32].

In addition to potential facilitation of MDRO acquisition or spread, antimicrobial overuse may also have negative impacts on the diversity, composition, or function of the gut microbiota, which may already be vulnerable in elderly populations. Healthy younger adults tend to have a fecal microbiome characterized by relatively high diversity of species and populated primarily by members of the phyla *Bacteroidetes* and *Firmicutes*, largely obligate anaerobes which exist in homeostasis with the intestinal epithelium[33–37]. However, it has been found that during senescence, the gut tends to have higher levels of *Bacteroidetes* and *Proteobacteria* and harbors higher levels of facultative aerobes and potential pathobionts, including *Enterobacterales* such as *E. coli* [36, 38–45]. These changes become more pronounced as aging progresses, and several studies have indicated that age-related alterations to the gut microbiota are relatively minor in septuagenarians, but become more pronounced over time and are clear in centenarians and supercentenarians[39, 43, 46–48]. This is likely due to a number of factors, including the decline of immune function, onset of age-related diseases (including metabolic disorders), changes to diet and mobility, and the increased likelihood of medication utilization and/or hospitalization[42, 49]. However, lifestyle of elderly adults has an important impact, as research suggests that community-resident elderly subjects have a distinct and more diverse microbiome compared with those of their hospitalized or institutionalized peers, which was suggested to be at least in part due to nutritional differences[49, 50]. Furthermore, reduced microbiome diversity has been associated with increased frailty of elderly subjects[49, 51]. Accordingly, given that the microbiomes of institutionalized elderly patients are perhaps already at risk, understanding the impacts of antimicrobial use and MDRO acquisition on this population is of importance.

We analyzed the gut microbiomes of eleven subjects from SPREAD to examine the impact of antimicrobial use on the gut microbiota composition, function, and antimicrobial resistance gene (ARG) profile of elderly dementia patients. These subjects were chosen as they were the largest cohort who had received a single antimicrobial (levofloxacin) during the collection period, and we anticipated that this intervention could have an impact on the already-vulnerable microbiota of this elderly, institutionalized cohort. Levofloxacin is an antimicrobial of the fluoroquinolone class with high oral bioavailability[52–54] which has been found to reduce levels of Gram-negative aerobic bacteria – including *Proteobacteria* and particularly *Enterobacterales* – in the fecal microbiota[55–61], although fluoroquinolone resistance among this taxon has been spreading[62–69]. A maximum of five rectal swab samples, collected every three months, were taken from each subject, and both 16S rRNA amplicon and shotgun metagenomics sequencing were performed. We analyzed alpha and beta diversity, taxonomic composition, functional potential, and antimicrobial resistance gene profiles before and after administration of levofloxacin, but were unable to detect specific impact of levofloxacin on any of these measures. However, we did find an association between blooms of particular enteric species and ARG burden, including in samples where MDRO were not detected by culture, suggesting that certain pathobionts carrying high ARG burdens may frequently colonize this population and that metagenomics may allow detection of resistant bacteria not flagged by culture-based methods.

## RESULTS

### Overview of Subjects

Elderly patients in long-term care facilities, and particularly patients with advanced dementia, are frequently exposed to antimicrobials and are at high-risk of acquisition and carriage of MDRO[9-13, 18-20, 27-30, 32]. From within the SPREAD cohort, we selected the largest group of subjects who had been administered a single antimicrobial during their participation in the study. This gave us a group of eleven subjects who had been given the fluoroquinolone levofloxacin, one of the most commonly prescribed antimicrobials. We analyzed up to five rectal swabs, taken every three months over the course of a year, from these eleven subjects in the SPREAD cohort[31], using both 16S rRNA and shotgun metagenomics sequencing (Figure 1A). During their participation in the study, these subjects had received only a single course of levofloxacin (average course of eight days), which has previously been shown to decrease the proportion of the *Enterobacterales* order of *Proteobacteria*[55–61]. Of the eleven subjects, all but Subject I were female and all but Subject G were white. They ranged in age from 72 to 101, and six members of the cohort did not survive for the full year of the study (Additional Table 1). All but two subjects (C and G) resided in different nursing homes. Overall, there were 38 samples for metagenomics sequencing (Additional Table 2). Culture-based methods indicated that four of the eleven subjects acquired a MDRO during the study: Subject A acquired methicillin-resistant *S. aureus* (MRSA) at the 12-month timepoint, Subject B acquired multidrug-resistant *E. coli* at the 3-month timepoint, and Subjects C and D both acquired multidrug-resistant *P. mirabilis* at the 3-month timepoint (Additional Table 1).

**Figure 1:**
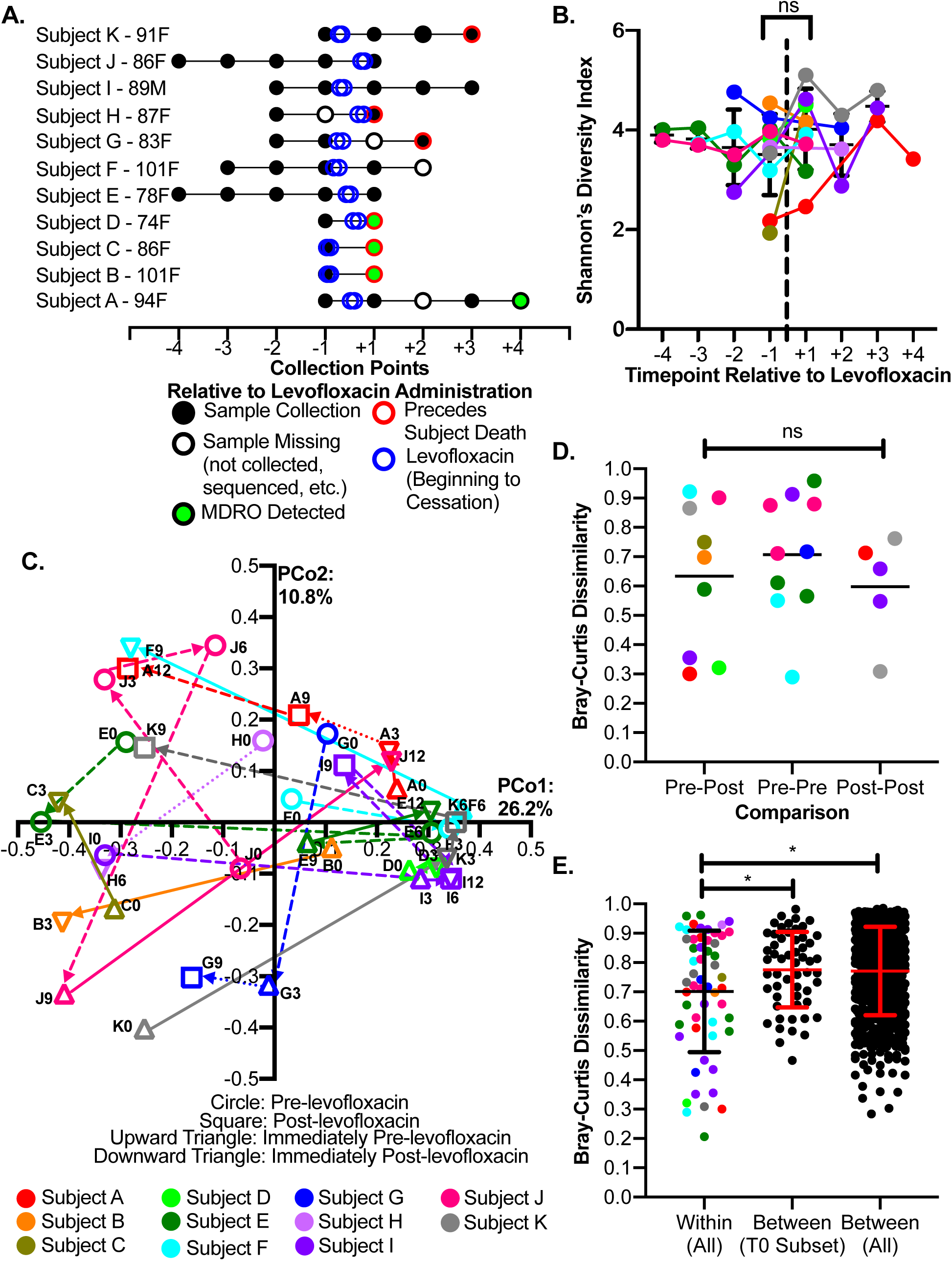
Subject Overview and Diversity Metrics. (A) Metagenomics sequencing was performed on longitudinal samples from eleven subjects from SPREAD who had received a single course of levofloxacin during their participation in the study. Points represent collection of samples, at intervals of approximately 3 months, relative to administration of levofloxacin. (B) Shannon diversity over time of all subjects. The dashed line indicates administration of levofloxacin. p = 0.175 for immediately pre-levofloxacin vs. immediately post-levofloxacin samples and p = 0.1006 for all pre-levofloxacin vs. all post-levofloxacin samples; Mann-Whitney test. (C) PCoA analysis of Bray-Curtis Dissimilarity. Solid arrows connect immediately pre- with immediately post-levofloxacin samples, dashed arrows connect other sequential samples, and dotted arrows connect samples where an intermediate sample is missing. (D) Within-subjects Bray-Curtis Dissimilarity of sequential samples. p = 0.6248 between pre-levofloxacin samples, post-levofloxacin samples, or pre-post levofloxacin samples; ANOVA). (E) Overall within-subjects, T0 between-subjects, and overall between-subjects Bray-Curtis Dissimilarity. p = 0.0262 for overall within-subjects vs. T0 between-subjects and p = 0.0175 for overall within-subjects vs. overall between-subjects; t-test with Welch’s correction.

### Alpha and Beta Diversity Metrics

Before focusing on antimicrobial resistance, we first wanted to assess the composition of the community throughout the longitudinal timeframe. We initially used the metagenomic sequencing data to compare the alpha diversity, or the diversity within each sample, of samples collected before and after levofloxacin administration. According to Shannon’s Diversity Index, which incorporates both richness and evenness of samples, there was no significant difference between the pre- and post-levofloxacin samples (Figure 1B). Furthermore, the alpha diversity was variable over time even within the same subject, and there was no clear trend of recovery in alpha diversity after antibiotic cessation. This suggests a degree of temporal instability, in which the richness and/or evenness of the samples varies changes over time.

We then examined beta diversity, or the diversity between samples. We utilized the Bray-Curtis Dissimilarity metric, which considers the identity and abundance of taxa shared between samples. Plotting this metric in a principal coordinate analysis (PCoA) revealed no apparent pattern of clustering based on either subject or sample collection point relative to levofloxacin, and in fact, samples from the same subject were often located quite distantly from one another (Figure 1C). We then compared the within-subjects dissimilarity of sequential samples within a subject when both were pre-levofloxacin, both were post-levofloxacin, or one sample was pre- and one was post-levofloxacin; there was no significant difference between any of the groups (Figure 1D), suggesting that levofloxacin was not associated with community disruption. Furthermore, while within-subject dissimilarity was lower than between-subjects dissimilarity, the effect size was low (0.7013 vs. 0.7712, respectively; Figure 1E).

### Taxonomic Composition

We utilized Kraken2 in conjunction the with Bayesian Reestimation of Abundance with KrakEN2 (Bracken2) pipeline to assign taxonomy to our metagenomic sequencing samples[70, 71]. Corresponding to the high between-subjects beta-diversity, the taxonomic composition of the gut microbiome varied significantly between subjects. As is typical for the human gut microbiome, most bacteria belonged to the five major phyla of *Firmicutes*, *Bacteroidetes*, *Proteobacteria*, *Actinobacteria*, and *Verrucomicrobia*. However, consistent with the high within-subjects beta diversity, the dominant phyla varied greatly even between samples from the same subject (Additional Figure 1); for example, the most abundant phylum in Subject E was *Bacteroidetes* at two timepoints, *Proteobacteria* at two timepoints, and *Firmicutes* at one (Additional Figure 1F). Overall, the most abundant phylum was *Actinobacteria* in three samples, *Bacteroidetes* in seventeen samples, *Firmicutes* in seven samples, and *Proteobacteria* in eleven samples (Additional Figure 1A-L); averaging across all samples, *Bacteroidetes* was highest at 34.2%, followed by *Proteobacteria* (26.9%), *Firmicutes* (23.3%), and *Actinobacteria* (11.2%) (Additional Figure 1A). Qualitatively, many of the samples from this population represent highly divergent and dysbiotic microbiomes compared with what is typically seen with younger subjects, in which *Proteobacteria* in particular make up a much smaller proportion of the microbiome than in these elderly dementia subjects[33].

The genus- and species-level taxonomic composition was also variable. Blooms of potential pathogens[72], including *Campylobacter ureolyticus*[73]*, Corynebacterium urealyticum*[74], *Enterococcus faecalis*[75, 76], *Escherichia coli* [77, 78], *Oligella urethralis*[79–82], *Proteus mirabilis*[83, 84], *Providencia stuartii*[85, 86], *Pseudomonas aeruginosa*[87, 88], *Staphylococcus aureus*[89–91], and *Staphylococcus haemolyticus*[92–94], were fairly common, both before and after levofloxacin administration (Figure 2A, Additional Figure 2). Across subjects, even baseline samples varied in composition, as expected from beta-diversity analysis. Averaging across all samples, the single most-abundant species was *E. coli*, further supporting the qualitatively dysbiotic nature of the gut microbiome of this cohort (Figure 2A). Despite the high proportion of members of *Enterobacterales* in this cohort, Linear Discriminant Analysis Effect Size (LEfSe) analysis[95] did not reveal biomarkers for pre- or post-levofloxacin samples at the phylum, genus, or species level. Full data on taxonomic composition at the phylum and species levels can be found in Additional Data 1.

**Figure 2:**
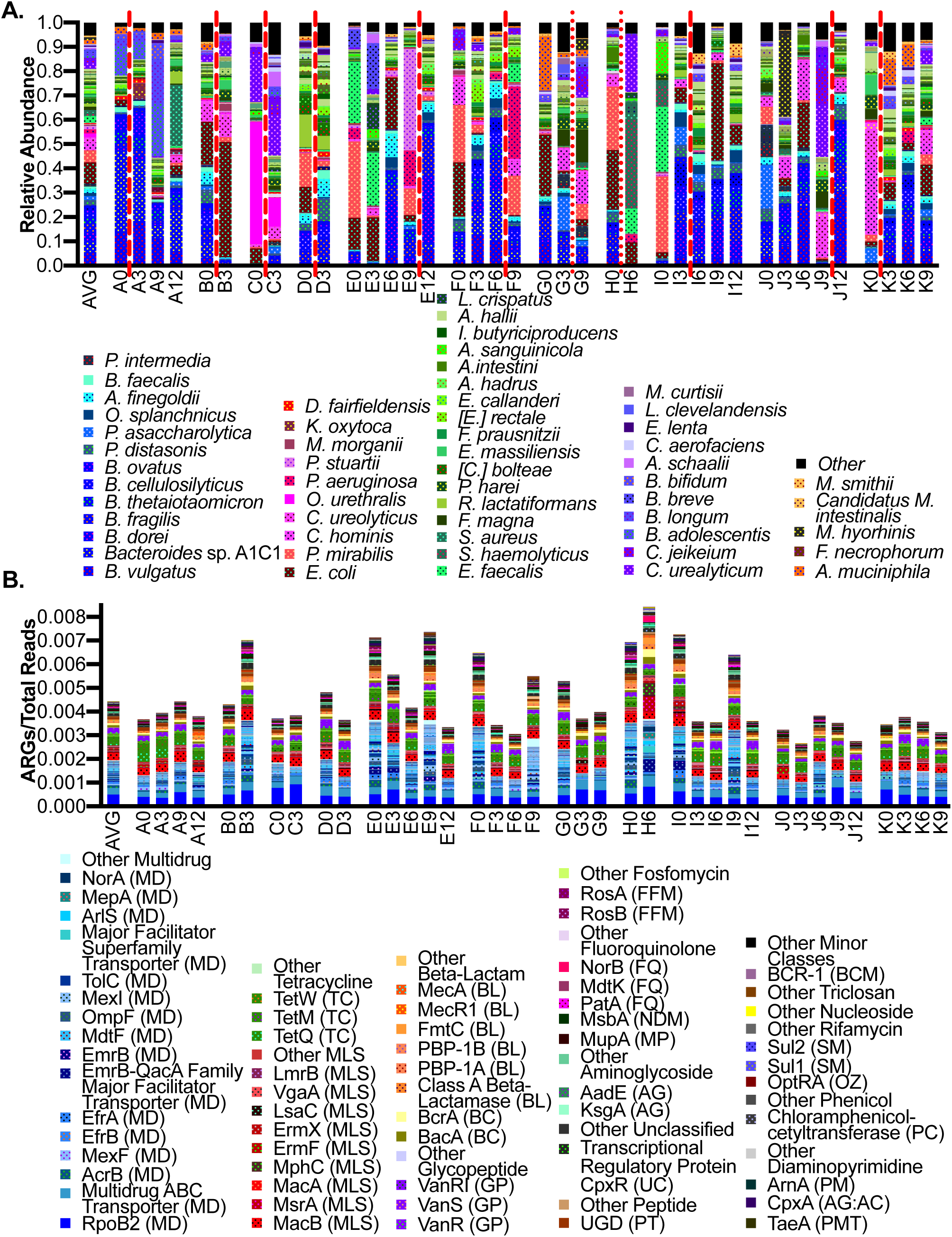
Relative Abundances of Species and Antimicrobial Resistance Genes. (A) Relative abundance of the most-abundant species across all samples, with all other species grouped in the “other” category. Species are grouped by genus and phylum, and are ranked within those levels by average relative abundance across all samples. Broad color categories distinguish phylum (*Proteobacteria* are red, *Bacteroidetes* are blue, *Firmicutes* are green, and *Actinobacteria* are purple), while different species of the same genus are given the same specific background color. Red lines indicate levofloxacin administration; dashed lines indicate usage between consecutive timepoints, while dotted lines indicate usage where the immediately post-levofloxacin sample is missing (B) Relative abundance of the most-abundant antimicrobial resistance genes (ARGs) across all samples. Specific ARGs are grouped by the class of antimicrobials they provide resistance to. Broad color categories distinguish class (Multidrug RGs are blue, MLS RGs are red, etc.), while related gene categories (ex: the *mec* operon or *mex* efflux proteins) are given the same specific background color. All ARGs were normalized to the total number of reads.

As we had access to full 16S rRNA and shotgun metagenomics data for our samples, we compared their taxonomic identifications at the genus level. The two methods of analysis were generally consistent, and blooms of prominent genera (including *Escherichia, Proteus, Enterococcus, Providencia, Staphylococcus*, and *Bacteroides*) were generally detected by both analysis pipelines (Additional Figure 3A). Metagenomics analysis was unsurprisingly able to detect more distinct genera, and of the genera that were called by both pipelines, LEfSe analysis revealed biases in both methods. For example, metagenomics analysis by Kraken2 and Bracken2 detected higher levels of *Bacteroides*, while 16S rRNA analysis with Quantitative Insights Into Microbial Ecology 2 (QIIME2)[96] detected higher levels of *Ruminiclostridium*. Full data on taxonomic abundances at the genus level can be found in Additional Data 1 for metagenomics and Additional Data 2 for 16S rRNA.

**Figure 3:**
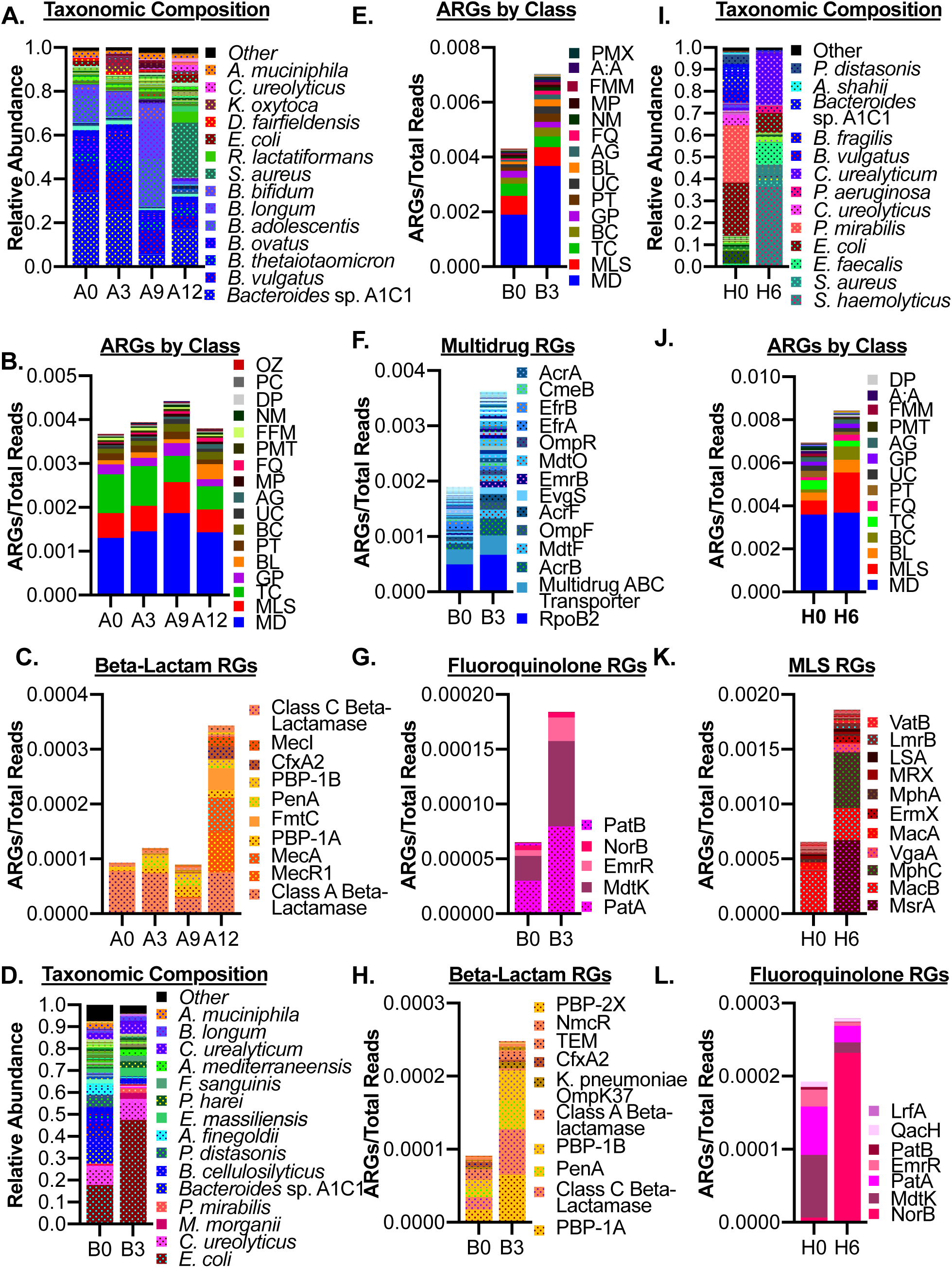
Antimicrobial Resistance Gene Profiles Reflect Taxonomic Observations. (A) Relative abundance of species in Subject A, showing a bloom in *S. aureus* at T12. (B) Relative abundance of ARG classes in Subject A, showing an expansion in beta-lactam resistance genes at T12. (C) Relative abundance of beta-lactam resistance genes in Subject A, showing increases in the *mecA/mecI/mecRI* operon at T12. (D) Relative abundance of species in Subject B, showing a bloom in *E. coli* at T3. (E) Relative abundance of ARG classes in Subject B, showing an expansion in multidrug, beta-lactam, and fluoroquinolone resistance genes at T3. (F) Relative abundance of multidrug resistance genes in Subject B, showing increases in various ARGs at T3. (G) Relative abundance of fluoroquinolone resistance genes in Subject B, showing increases in genes including *patA* and *mdtK* at T3. (H) Relative abundance of beta-lactam resistance genes in Subject B, showing increases in genes including penicillin-binding proteins and class C beta-lactamase at T3. (I) Relative abundance of species in Subject H, showing a bloom in *S. haemolyticus* at T6. (J) Relative abundance of ARG classes in Subject H, showing increases in MLS and fluoroquinolone resistance genes. (K) Relative abundance of MLS resistance genes in Subject H, showing an increase in staphylococcal resistance gene *msrA* and others at T6. (L) Relative abundance of fluoroquinolone resistance genes in Subject H, showing an increase in staphylococcal resistance gene *norB* and others at T6.

### Functional Potential

We used the Human Microbiome Project Unified Metabolic Analysis Network 2 (HUMAnN2) pipeline[97] to analyze the genetic content of the metagenomic samples. We utilized LEfSe to compare community function at the Kyoto Encyclopedia of Genes and Genomes (KEGG) ortholog, Gene Ontology (GO) term, and MetaCyc pathway levels. As in the taxonomic analysis, there were no significant biomarkers of either pre-or post-levofloxacin administration samples. However, while the taxonomic profile of the samples varied greatly, the functional capacity of the samples was fairly consistent across samples (Additional Figure 4). Full data on functional potential can be found in Additional Data 3.

**Figure 4:**
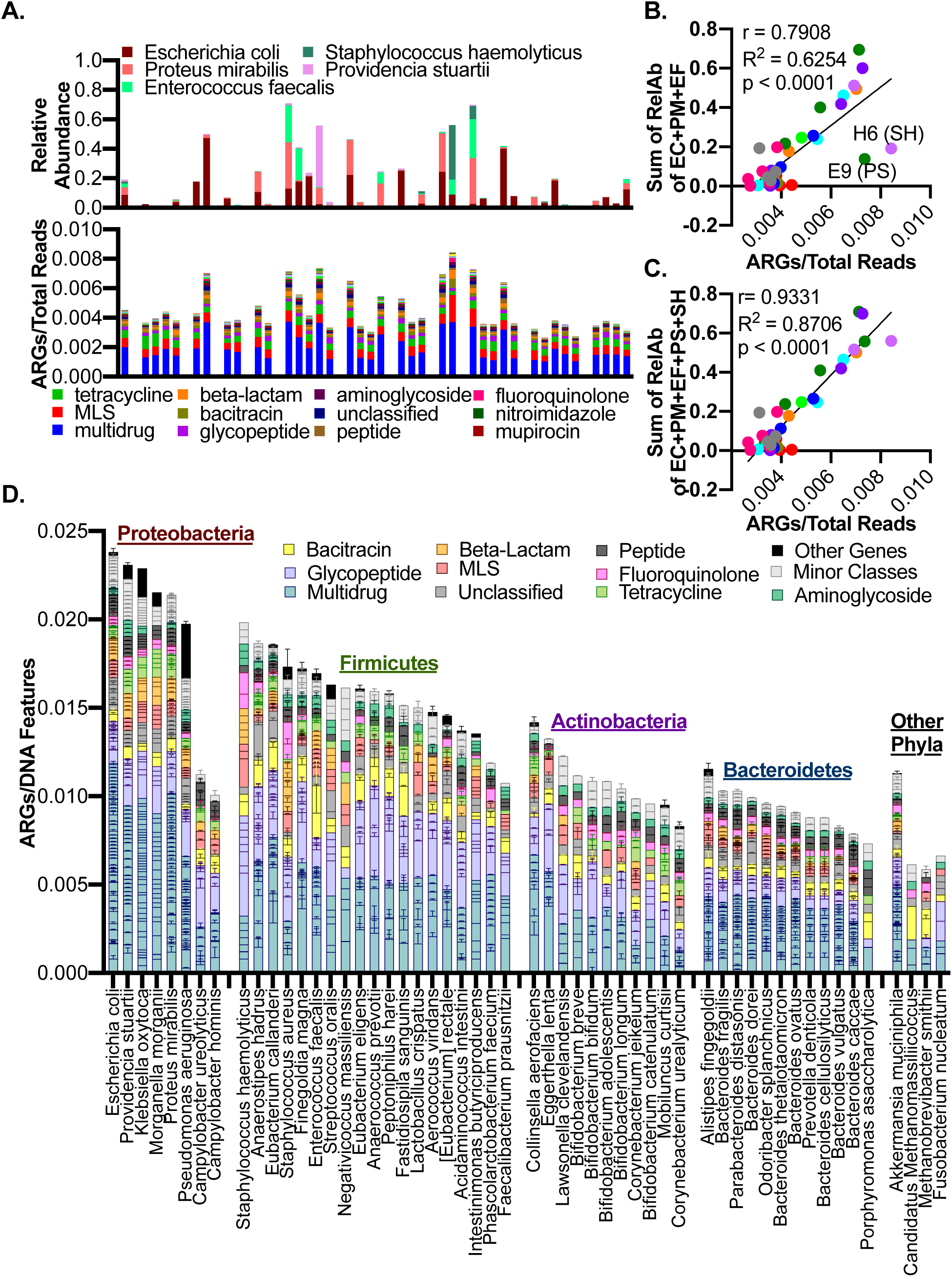
Relationship of ARG Levels to the Relative Abundance of Specific Pathobionts. (A) Correspondence between the relative abundances of key species of interest (*E. coli*, *P. mirabilis*, *E. faecalis*, *P. stuartii*, and *S. haemolyticus*) and total ARG density in each sample. (B) Correlation between the sum of the relative abundances of *E. coli*, *P. mirabilis*, and *E. faecalis* and the total ARG density in each sample (r = 0.791, R^2^ = 0.6254, p <0.0001; Pearson’s correlation). (C) Correlation between the sum of the relative abundances of *E. coli*, *P. mirabilis*, *E. faecalis*, *P. stuartii*, and *S. haemolyticus* and the total ARG density in each sample (r = 0.933, R^2^ = 0.8706, p < 0.0001; Pearson’s correlation). (D) Average ARG density in bins of species across all samples in which we were able to construct a bin for that species. Specific genes are grouped and colored by their ARG class, and bins are grouped by phylum and ranked by their total average ARG density within that phylum.

### Antimicrobial Resistance Gene Profile

We used the DeepARG machine-learning program[98] to detect resistance genes in the metagenomic samples. Across all samples, the most abundant class of ARG was “multidrug”, followed by “macrolide-lincosamide-streptogramin” (MLS), and “tetracycline”. The most common specific gene detected was the multidrug resistance *rpoB2* variant of the RNA polymerase beta subunit, followed by the MLS resistance gene *macB* and a multidrug ABC transporter (Figure 2B). LEfSe analysis revealed no ARG biomarkers of either pre- or post-levofloxacin samples. Full data on ARG composition can be found in Additional Data 4.

However, we were able to detect changes in specific ARG classes and genes that corresponded with the detection of antimicrobial-resistant organisms in two subjects. Subject A acquired MRSA at the 12-month timepoint, and a bloom of this species to 25.0% could be detected in the metagenomic taxonomic data (Figure 3A, Additional Figure 2B). While the overall level of ARGs did not notably increase at this timepoint, there was a clear expansion in beta-lactam resistance genes (Figures 3B, Additional Figure 5B), including the *mecA*/*mecR1*/*mecI* operon, which regulates expression of the low-affinity penicillin-binding protein *mecA* (PBP-2A)[99–102] (Figure 3C). This operon is characteristic of MRSA strains[99–102], supporting the culture-based classification of this *S. aureus* isolate as MRSA.

**Figure 5:**
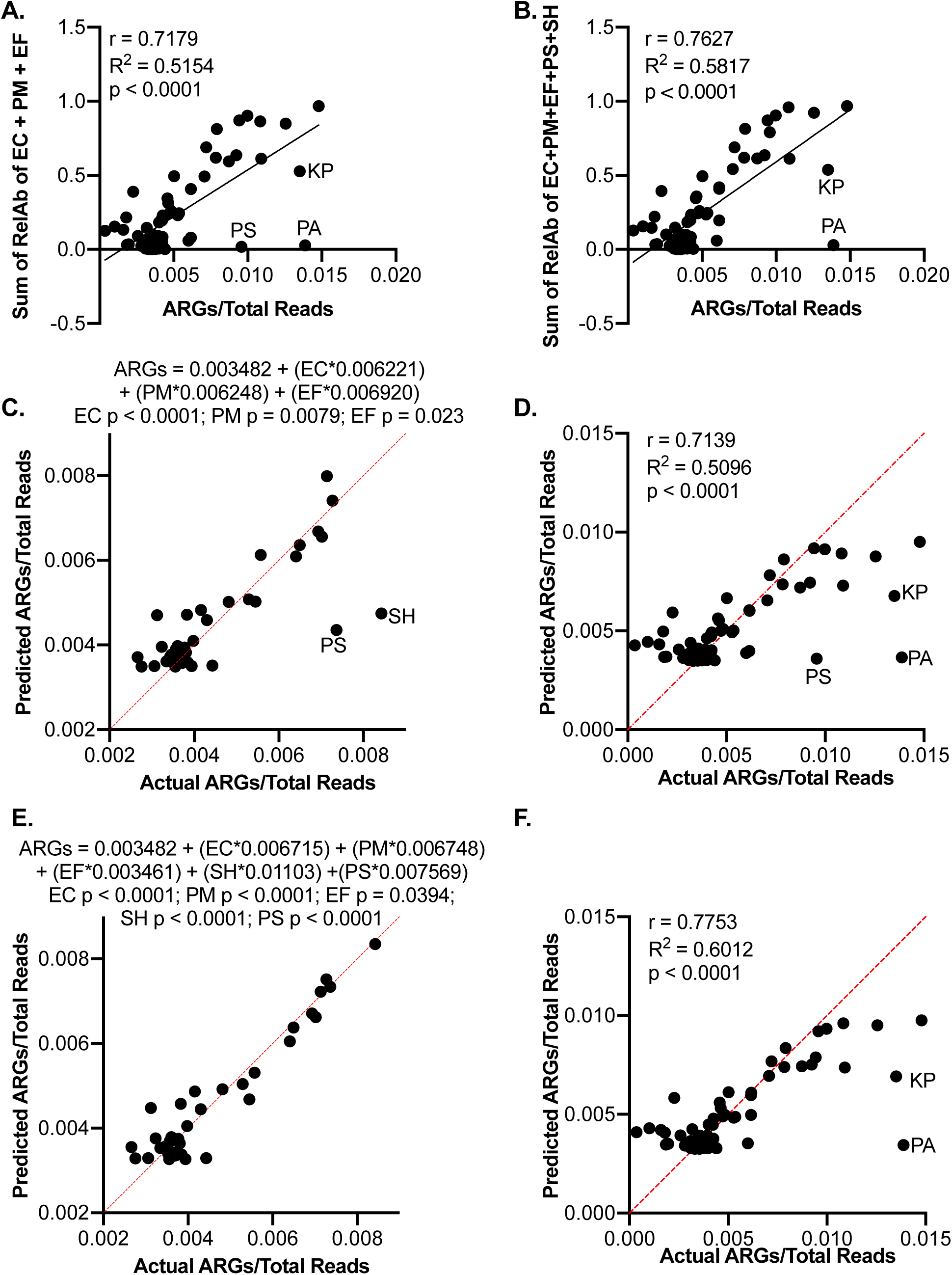
Prediction of ARG Density From Relative Abundance of Specific Pathobionts. (A) Correlation between the sum of the relative abundances of *E. coli*, *P. mirabilis*, and *E. faecalis* and the total ARG density in each sample in the test dataset (r = 0.7139, r^2^ = 0.5096, p<0.0001; Pearson’s correlation). (B) Correlation between the sum of the relative abundances of *E. coli*, *P. mirabilis*, *E. faecalis*, *P. stuartii*, and *S. haemolyticus* and the total ARG density in each sample in the test dataset (r = 0.7753, r^2^ = 0.6012, p<0.0001; Pearson’s correlation). (C) Multiple linear regression of relative abundances of *E. coli*, *P. mirabilis*, and *E. faecalis* to ARG density in samples in the levofloxacin dataset (38 samples). (D) Correlation between the predicted ARG density and actual ARG density in the test dataset (67 samples) based on the relative abundances of *E. coli*, *P. mirabilis*, and *E. faecalis*. (E) Multiple linear regression of relative abundances of *E. coli*, *P. mirabilis*, *E. faecalis*, *P. stuartii*, and *S. haemolyticus* to ARG density in samples in the levofloxacin dataset (38 samples). (F) Correlation between the predicted ARG density and actual ARG density in the test dataset (67 samples) based on the relative abundances of *E. coli*, *P. mirabilis*, *E. faecalis*, *P. stuartii*, and *S. haemolyticus*.

Similarly, Subject B acquired multidrug-resistant *E. coli* (resistant to the beta-lactams ampicillin/sulbactam, cefazolin, ceftazidime, and ceftriaxone and to the fluoroquinolone ciprofloxacin) at the three-month timepoint, and the proportion of this species expanded to 47.3% of the population in the corresponding sample (Figure 3D, Additional Figure 2C). Accordingly, this sample showed a notable increase in the relative abundance of ARGs, which was in large part driven by an increase in a number of multidrug resistance genes (Figure 3E); there was also a clear increase in several beta-lactam resistance genes, including the low-affinity penicillin-binding protein genes *PBP-1A, PBP-1B,* and *penA* (*PBP2*) as well as class C beta-lactamase genes[103–108], and several fluoroquinolone resistance genes, including the transporters *patA* and *mdtK* [109–113] (Figure 3F-G).

Despite the acquisition of multidrug-resistant *P. mirabilis* at the three-month timepoint in Subjects C and D, there was no corresponding increase in ARGs. ARG levels stayed approximately the same in Subject C (0.372% at baseline and 0.384% at three months) and decreased in Subject D from 0.482% at baseline to 0.364% at the three-month timepoint (Figures 2B, Additional Figure 5D-E). However, this corresponds to the taxonomic data; levels of *P. mirabilis* were low and stable in Subject C (0.55% at baseline and 0.61% three months later), and while *P. mirabilis* made up 13.8% of the population at baseline in Subject D, it underwent a reduction to 2.3% at the three-month timepoint (Figures 2A, Additional Figure 2D-E). Taken together, this data indicates that our metagenomics pipeline can detect blooms of AMR pathogens and that the corresponding change in ARG levels aligns with culture-based detection of MDROs. At the same time, metagenomic analysis of some samples found blooms of pathogens and ARGs that were not associated with culture-based MDRO detection.

### Attribution of ARG Density to Specific Species

While total ARG density within samples did not vary by levofloxacin administration, there was significant variability between samples. In fact, most samples had similar baseline levels of ARGs of 0.3% to 0.4% of the total reads, while only a few samples rose above this value to between 0.6% and 0.8%. Close inspection of the taxonomic composition of the samples revealed that samples with higher levels of ARGs tended to have blooms of one or more of the *Proteobacteria* species *E. coli* and *P. mirabilis* and the *Firmicutes* species *E. faecalis*, strains of which are common pathobionts[114–120] (Figure 4A). Confirming this association, correlation analysis between ARG levels and the sum of the relative abundances of these three species showed a strong and significant positive correlation (r = 0.791, R^2^ = 0.6254, p <0.0001, Pearson’s correlation; Figure 4B). This suggests that in samples with higher-than-baseline ARG levels, ARG abundance is being driven by high relative abundance of these three species.

However, there were two notable exceptions: Samples E9 and H6 had high levels of ARGs without corresponding blooms of these three species. However, *P. stuartii* bloomed to 41.9% relative abundance in Sample E9 and *S. haemolyticus* bloomed to 36.9% in Sample H6 (Figure 2A, Additional Figure 2F&I). Both species have long been associated with AMR phenotypes[93, 94, 121–127] and were not found at high levels in other samples, but could explain the higher ARG abundance in these samples (Figure 4A). Supporting this possibility, an examination of the ARGs in Sample H6 showed a distinct profile relative to other samples, with high levels of staphylococcal resistance genes including fluoroquinolone resistance gene *norB* and macrolide-streptogramin resistance gene *msrA* [128–131](Figures 2B, 3I-L). Accordingly, addition of *P. stuartii* and *S. haemolyticus* abundances to the analysis resulted in a stronger correlation (r = 0.933, R^2^ = 0.8706, p < 0.0001, Pearson’s correlation; Figure 4C).

To more rigorously examine the relationship between the species of interest and ARG levels, we performed metagenomic assembly and binning to compare the levels of ARGs in these organisms to levels in other common and abundant species, including likely commensals and potential pathogens (Figure 4D). Specifically, we analyzed bins that passed various quality controls (Additional Table 3) and corresponded to species identified by Kraken2/Bracken2 to make up greater than 0.1% of their source samples (Additional Table 4).

As anticipated, we found that the levels of ARGs in bins from *E. coli* and *P. mirabilis* were consistently high compared to other species analyzed. In fact, *E. coli* had the highest average ARG density of any species analyzed, while *P. mirabilis* was the fifth-highest. Notably, the ARG composition of the bins of these species from samples in which MDROs were detected (B3, C3, and D3) did not appear to be different from those of other samples (Additional Figure 6A-B), although it is possible that some resistance genes were carried on plasmids that were not assembled into genomes. *P. stuartii* had the second-highest average ARG density, reflecting the expansion of ARGs detected in sample E9, where this species bloomed to 41.9% of the population. The third and fourth positions were taken by the single bins constructed for *Klebsiella oxytoca* and *Morganella morganii*, other *Proteobacteria* with pathogenic potential[132–134]. *P. aeruginosa* bins rounded out the top six, with similar levels to the other top species. However, as *K. oxytoca* and *M. morganii* were never present at greater than 3% and *P. aeruginosa* bloomed in only two samples, they did not significantly contribute to overall ARG density in the cohort. Importantly, high ARG density was not a universal feature of *Proteobacteria*, or even of pathogenic *Proteobacteria*; bins constructed for the *Campylobacter* species *C. hominis* and *C. ureolyticus* had universally low ARG levels. Additionally, while we could not construct a high-quality bin for *O. urethralis*, the low ARG densities in the samples in which this species bloomed (C0 and C3) suggests that it also has low genomic ARG content. This suggests that high ARG density among the *Proteobacteria* analyzed was restricted to the *Gammaproteobacteria* class, primarily of the order *Enterobacterales* but also including *Pseudomonadales*.

We were only able to construct two good-quality bins for *E. faecalis*, which varied in their ARG levels, particularly on the basis of bacitracin resistance. On average, while the two bins did not have ARG levels as high as the *Proteobacteria* of interest, they did rank among the highest of the *Firmicutes* bins tested. We were also able to create a single bin for *S. haemolyticus* from Sample H6 in which it made up 36.9% of the population. This bin had an ARG density higher than the average for any other non-*Proteobacteria* species, supporting its role in the high ARG levels found in the corresponding sample. As expected from the analysis of the total ARG population of that sample (Figure 3G), the staphylococcal resistance genes *norB* and *msrA* were found in this bin. We were also able to create two bins for *S. aureus*, including from sample A12 where MRSA was detected. The A12 bin contained the characteristic MRSA gene *mecA* while the H6 bin did not, suggesting that the *S. aureus* strain found in H6 was not MRSA (Figure S6C). In general, bins from the phyla *Actinobacteria* (including *Bifidobacterium* and *Corynebacterium* species) and *Bacteroidetes* (including *Bacteroides* and *Parabacteroides* species) had low ARG levels. Full data on the ARGs and classes found in species-level bins can be found in Additional Data 4.

### Prediction of ARG Density from Species Abundances

Our initial analysis only considered the eleven subjects for whom we had longitudinal metagenomics data due to their receiving levofloxacin. We also had access to a larger dataset: shotgun metagenomics had been performed on a further 67 samples for a related study. In this case, the data was not longitudinal and encompassed an array of antibiotic treatment conditions across 67 subjects, providing a diverse set of taxonomic and ARG data on which to test whether the relationship between *E. coli*, *P. mirabilis*, and *E. faecalis* and ARG density held true. As an initial test, we performed the same correlation analyses between species of interest and ARG levels as on the levofloxacin dataset, finding that both the simple and complex models showed significant correlation (r = 0.7179, r^2^ = 0.5154, p < 0.0001 and r = 0.7627, r2 = 0.5817, p<0.0001, respectively; Pearson’s correlation; Figure 5A-B). This provided initial support for the trend being present in the wider dataset.

We then created a multiple linear regression model to predict ARG density using the relative abundances (RA) of the three main species of interest in the initial levofloxacin dataset, with the following equation: (ARG density) = 0.003482 + 0.006221(*E. coli* RA) + 0.006248(*P. mirabilis* RA) + 0.006920(*E. faecalis* RA) (Figure 5B). We then used this equation to predict the ARG density in the larger metagenomics dataset and found that it was able to accurately predict the true ARG level of those samples, with predicted and actual values correlating significantly (r = 0.7139, r^2^ = 0.5096, p<0.0001; Pearson’s correlation; Figure 5C). As before, there were a few notable outliers with higher ARG levels than predicted by the model; those three samples contained high levels of *P. stuartii*, *P. aeruginosa*, or *Klebsiella pneumoniae* This maps well to the fact that we observed high levels of ARGs in bins constructed from *P. stuartii*, *P. aeruginosa*, and the related species *K. oxytoca* (Figure 4D).

We also created a multiple linear regression model that incorporated the relative abundances of *P. stuartii* and *S. haemolyticus*, which caused outliers from the original species-ARG correlation: (ARG density) = 0.003253 + 0.006715(*E. coli* RA) + 0.006748(*P. mirabilis* RA) + 0.003461(*E. faecalis* RA) + 0.01123(*S. haemolyticus* RA) + 0.007569(*P. stuartii* RA) (Figure 5E). As before, we tested this equation against the larger dataset, and found that it slightly increased the accuracy of the predictions; specifically, it eliminated the outlier which had high *P. stuartii* levels and slightly improved the correlation between predicted and actual ARG levels (r = 0.7753, r^2^ = 0.6012, p<0.0001; Pearson’s correlation; Figure 5F). However, the simpler model is more broadly applicable, as blooms of *P. stuartii* and *S. haemolyticus* are relatively uncommon.

Similarly, while *Klebsiella* spp. and *P. aeruginosa* may also contribute to high ARG density in samples, they do not bloom as commonly in this cohort as the core predictive species of *E. coli*, *P. mirabilis*, and *E. faecalis*.

These results indicate that in this population, levels of only a few key species could predict the majority of ARG abundance beyond background levels. Both the core predictive species (*E. coli*, *P. mirabilis*, *E. faecalis*) and others that are associated with high ARG levels in samples (*P. stuartii*, *S. haemolyticus*, *P. aeruginosa*, *Klebsiella* spp.) are pathogens and/or pathobionts. Monitoring levels of these species may be helpful in elderly, institutionalized populations, as these patients may be vulnerable to developing or transmitting AMR infections from high-level carriage of these species.

## DISCUSSION

Overall, we found that the microbial composition of the gut microbiome of elderly patients with advanced dementia was quite variable, both between subjects and over time within the same subject. Even in the absence of antimicrobial treatment, there was notable fluctuation in the abundance of a number of species, including pathobionts such as *E. coli*, *P. mirabilis*, and *E. faecalis*. When comparing the taxonomic composition, functional potential, and resistome of pre- and post-levofloxacin samples, we did not observe any significant differences. One potential reason for this finding is that oral levofloxacin is well-absorbed by the host, with greater than 99% bioavailability[52, 53, 135–137], and therefore may not be directly available to the luminal microbiota of the lower gastrointestinal tract at high levels. Furthermore, other studies have suggested that levofloxacin has a relatively minor impact on the gut microbiome, primarily reducing levels of *Enterobacterales*[55–61], and it may be less-associated with *Clostridiodes difficile*-associated diarrhea outbreaks than other antimicrobials, including other fluoroquinolones[138].

Additionally, in this cohort, levofloxacin was typically administered at least two weeks prior to collected timepoints, potentially allowing sufficient time for the microbiome to recover from or shift away from its immediately post-antibiotic state. Furthermore, the impacts of levofloxacin on the gut microbiome may be dependent upon the initial state upon administration. If the microbiome is initially relatively diverse and healthy, antibiotic administration may be disruptive and allow blooms of atypical dominant species such as members of *Proteobacteria*; such an occurrence might be observed in Subject F, where a diverse *Bacteroides*-dominated microbiome was overtaken by several *Enterobacterales* after levofloxacin treatment (Additional Figure 2G). Alternatively, if the microbiome is initially dominated by one or more pathogens, antimicrobial administration may correct such blooms and allow for the restoration of a diverse community, as might have occurred in Subject E as a *P. stuartii* bloom was eliminated (Additional Figure 2F).

Finally, since the pre-existing temporal instability of this community was high, levofloxacin-related changes may not be detectable through the noise of this cohort’s general microbiome instability. In contrast to our observations, studies in adults have generally found that the within-subjects dissimilarity is much lower than between-subjects dissimilarity, in line with the fact that the gut microbiome tends to be relatively stable within the same subject over time – including in an elderly cohort[33, 34, 41, 139, 140]. This suggests that the gut microbiomes of the subjects in this study were less stable than that of other cohorts, potentially suggesting that this institutionalized population with advanced functional impairment is more prone to infections or has weaker immune systems than young healthy adults or even community-resident elderly adults.

Interestingly, despite the taxonomic variability, the functional composition of the cohort was relatively similar across samples and subjects. This is in line with previous studies of the human gut microbiome, which suggest that variable taxa can fill the same functional niches, resulting in a more similar functional composition across individuals despite inter-individual differences in the taxonomic composition[33, 35, 141].

As all of the subjects had been given an antibiotic, we were particularly interested in the antibiotic resistance profile of the subjects before and after levofloxacin administration. However, as observed in the taxonomic and functional data, there was no apparent association of any ARG genes or classes with either pre- or post-levofloxacin status. This may be due to the fact that levofloxacin did not have any specific impacts on the resistome of this cohort, or due to the factors that may have concealed any impacts of levofloxacin, as discussed above. However, we were particularly intrigued by the finding that ARG density in a particular sample could be linked to the abundance of a few key species. *E. coli*, *P. mirabilis*, and *E. faecalis* are all pathobionts that are often found at low levels in a healthy microbiome, but bloomed frequently at various timepoints across a majority of our subjects. All three species can cause severe illness, have been previously observed to colonize nursing home residents, and include well-known multidrug-resistant strains[11-13, 28, 30, 32, 68, 75, 114-120, 142]. In fact, three of the subjects (B, C, and D) are known to have acquired multidrug-resistant strains of *E. coli* and *P. mirabilis* during the study. However, we observed an association between these three species and ARG levels across the entire sample set (Figure 4B), and the ARG composition of the bins of *E. coli* and *P. mirabilis* from samples where MDROs were detected were not distinct from their other bins (Additional Figure 6A-B). This suggests that metagenomic sequencing may allow the identification of antimicrobial-resistant organisms that escape detection via culture-based techniques, although it is also possible that the multidrug-resistant isolates contained ARG-carrying plasmids that were not captured by our assembly and binning strategy.

A major implication of this finding is that metagenomic analysis could be a particularly useful tool to track antimicrobial resistance in institutions like nursing homes and hospitals, particularly with the capability to construct contigs and bins that allow examination of specific genomes. In this case, it has allowed us to connect the high levels of ARGs in certain samples with correspondingly high levels of specific pathobionts, which had high proportions of ARGs within their genomes even in samples where MDROs were not detected. In a vulnerable population already prone to infections and carriage of MDROs, metagenomics could be a useful surveillance tool to assess the prevalence or transmission of ARGs in long-term care facilities.

Importantly, all of the subjects in our study were institutionalized in nursing homes, and there exists significant potential for transfer of bacteria between patients. As all but two of our subjects (C and G) lived in different homes, we could not directly examine this possibility ourselves, but it is possible that the high abundance of pathobionts and/or ARGs in our cohort is related to the spread of isolates within nursing homes. This also raises the possibility that we would not find a similar association between pathobionts and ARG levels in a healthy or community-based elderly cohort, who might be less likely to harbor or transmit such high levels of these bacteria. However, if an association between particular “sentinel” species and ARGs holds true in other elderly institutionalized populations, qPCR detection of the loads of these such pathobionts may allow for prediction of resistant bacterial outbreaks before they occur.

In addition to the increased potential for spread of resistant strains through institutions, there are some other potential explanations for the association between ARGs and these particular species. In particular, all of the species that we found to be associated with ARG density are potential human pathogens, can be grown *in vitro*, and have been previously associated with AMR phenotypes. ARGs, as well as mobile genetic elements carrying them, from these species may be better-studied than those from organisms less likely to pose a threat to human health, including gut commensals. If ARGs from these organisms are well-represented in databases, it could potentially bias analyses based on these databases toward detecting pathogen-over commensal-associated ARGs. However, there has been significant work done on the resistome of the human commensal microbiome, including functional metagenomics to detect new ARGs. These have found that commensal anaerobes may serve as significant reservoirs of ARGs, and may in some cases contribute to the transfer of resistance to pathobionts[143–150]. Commensal carriage of antimicrobial resistance genes may correspond to the baseline level of 0.3-0.4% ARGs observed in samples without pathobiont dominance.

Some limitations to the findings of this study must be acknowledged. First, as for all database-based methodologies, we are limited by accuracy and completeness of those databases. While the human gut microbiome is fairly well-characterized, there may be so-called microbial dark matter that is not well-represented in the taxonomic database used for species identification. We also used a database composed of bacterial and archaeal genomes, excluding consideration of bacteriophage and microbial eukaryotes from our analyses. As mentioned, database representation is particularly relevant for our ARG analysis, as the genes in this database may be skewed towards easily-culturable and pathogenic source species, and our analysis may have missed ARGs found in commensal or unculturable gut species. Additionally, critics have noted that some genes found in ARG databases used have unclear links to resistance phenotypes, and may perform regulatory, efflux, or other functions not always related to antimicrobial resistance[143, 151].

Second, we were limited by the original SPREAD population, in which few subjects received only a single antimicrobial during the course of the study; this makes it difficult to say whether the temporal variability we observed was widespread in the cohort, although the fact that there were frequently high pathobiont levels observed in the larger metagenomics dataset we used to test our multiple linear regression suggests that this may be the case. Third, in this study we worked with rectal swabs, which are similar but not identical to fecal samples and may be susceptible to cross-contamination from urinary pathogens or skin flora, particularly in incontinent advanced dementia patients[152–155]. Fourth, metagenomic assembly has limitations. It cannot create bins of all species found in a given sample, genome reconstruction is based on the isolates present in the database used, and analysis of assembled genomes may exclude consideration of plasmids – which are often sources of ARGs. Finally, as we analyzed metagenomic data, we cannot comment on the actual antimicrobial resistance phenotypes of the communities or individual bacteria that we studied.

## CONCLUSIONS

The gut microbiome was highly variable both between and within subjects, with frequent blooms and reductions of bacterial species both before and after levofloxacin treatment. We did not observe a consistent impact of levofloxacin on specific taxa or functions, levels of antimicrobial resistance genes, or overall microbiome diversity in these subjects. However, while we could not link levofloxacin to antimicrobial resistance gene levels, there were a number of samples that had higher relative abundances of these genes. In our original metagenomics dataset, we were able to identify that levels of these genes could be linked to blooms of specific bacterial species, including *E. coli*, *P. mirabilis*, and *E. faecalis*. We were able to build a model to predict total ARG levels in a sample from the relative abundance of these species, and confirm the validity of this model in a larger metagenomics dataset from the rest of the SPREAD study, including subjects taking a range of antibiotics. Furthermore, use of metagenomic assembly and binning allowed us to confirm that our species of interest carry greater ARG densities than other abundant members of the microbiome, even in subjects where MDROs were not detected by culturing.

This demonstrates that there is a significant amount of information that can be obtained from metagenomic assembly and binning. With sufficient depth, powerful computational tools allow whole genomes to be assembled from short-read metagenomic sequencing, which permits interrogation of the likely features of species of interest in complex microbial communities. In our case, we were able to confirm the association between pathobiont blooms and ARG levels in the gut, showing that the genomes of pathobionts contained a greater proportion of ARGs than gut commensals such as *Bacteroides* and *Bifidobacterium* species. This suggests that while the commensal microbiota are known to serve as reservoirs of antimicrobial resistance, in this cohort blooms of pathobionts may serve as the driver of ARG levels in the gut microbiome. Given how frequently these blooms occurred, special attention should be paid to these species in dementia patients in long-term care facilities, a vulnerable group which is often immunocompromised, frequently administered medication including antimicrobials, and may carry MDRO at relatively high levels.

## Abbreviations

A:A: Aminoglycoside:Aminocoumarin
AG: Aminoglycoside
AMR: Antimicrobial Resistant
ARG: Antimicrobial Resistance Gene
BC: Bacitracin
BL: Beta-Lactam
Bracken2: Bayesian Reestimation of Abundance with KrakEN2
DP: Diaminopyrimidine
FFM: Fosfomycin
FMM: Fosmidomycin
FQ: Fluoroquinolone
GP: Glycopeptide
GO: Gene Ontology
HUMANn2: Human Microbiome Project Unified Metabolic Analysis Network 2
KEGG: Kyoto Encyclopedia of Genes and Genomes
LEfSe: Linear Discriminant Analysis Effect Size
MD: Multidrug
MDRO: Multidrug-Resistant Organism
MLS: Macrolide-Lincosamide-Streptogramin
MP: Mupirocin
MRSA: Methicillin-resistant *Staphylococcus aureus*
NM: Nitromidazole
OZ: Oxazolidinone
PATRIC: Pathosystems Resource Integration Center
PC: Phenicol
PMT: Pleuromutilin
PMX: Polymyxin
PT: Peptide
QIIME2: Quantitative Insights Into Microbial Ecology 2
TC: Tetracycline
SPREAD: Study of Pathogen Resistance and Exposure to Antimicrobials in Dementia
UC: Unclassified

## METHODS

### Sample Collection and Preparation

#### Subjects

Eleven subjects were chosen from the SPREAD cohort based on the following inclusion criteria: at least two consecutive rectal swabs were collected from the subject during the study, subjects had received a single oral course of levofloxacin during the study (average course of 8 days), and subjects received no other antimicrobials during the study or in the 3 months prior to the first sample collection. Of the 11 subjects, 10 were female and 10 were white, while ages ranged from 72 to 101. Five subjects lived through the entire sample collection period, while the other six passed away at some point prior to the final collection; between this attrition, one sample that was not collected, and three samples that were not well-sequenced, we had a total of 38 usable metagenomic samples (Figure 1A; Additional Tables S1-2). All samples were collected under SPREAD, which was approved by the Institutional Review Board at Hebrew Life[31].

#### Sample Collection

Samples were collected by insertion of sterile double-tipped swabs (Starswab II; Starplex Scientific Inc., Ontario, Canada) into the anus of the subject. The first swab was used to identify multidrug-resistant organisms (including methicillin-resistant *S. aureus*, vancomycin-resistant enterococci, and multidrug-resistant Gram-negative organisms such as *E. coli*, *P. mirabilis*, *P. aeruginosa*, or *P. stuartii*) via culturing techniques as described previously[156]. The second swab was frozen in 20% glycerol at -80°C for DNA extraction and sequencing.

#### Sample Processing

Frozen rectal swabs were thawed and placed into 96-well plates, before extraction using the PowerSoil DNA Isolation Kit (MOBIO, West Carlsbad, CA) according to the manufacturer’s instructions. DNA concentrations were measured using a Nanodrop 1000 (Thermo Scientific, Waltham, MA) and extracted DNA was stored at -20°C until further use.

### 16SrRNA Amplicon Sequencing

#### Sequencing

The V4 hypervariable region of the 16S rRNA gene was amplified according to Earth Microbiome Project protocols. Amplification was performed using Illumina-adapted universal 16S primers 515F and 806R under the following conditions: 3 minutes at 94°C, 45 cycles of [45 seconds at 94°C, 60 seconds at 50°C, 90 seconds at 72°C], 10 minutes at 72°C. All reactions were prepared using 5 PRIME polymerase 1X HotMasterMix (5PRIME, Gaithersburg, MD) and run in triplicate to alleviate primer bias. Triplicates were pooled before cleaning with a PCR Purification Kit (Qiagen). These products were quantified using the Qubit dsDNA High Sensitivity Assay Kit (Invitrogen, Eugene, OR) and samples were pooled in equimolar amounts. Sequencing was performed using the Illumina MiSeq platform located at the New York University Langone Medical Center Genome Technology Core. Sequences can be found under the BioProject accession number PRJNA573963 (http://www.ncbi.nlm.nih.gov/bioproject/573963).

#### Data Processing

Data processing was performed using the QIIME2 (v 2019.1) pipeline[96]. The Divisive Amplicon Denoising Algorithm 2 (DADA2) method was used to quality-filter sequences and categorize amplicon sequence variants (ASVs)[157], and the SILVA (release 132) 99% identity V4 classifier was used to assign taxonomy to ASVs[158]. See Additional File 1 for more information. Taxonomic relative abundances were exported at the genus level for further analysis. Output data can be found in Additional Data 2.

### Shotgun Sequencing

#### Sample Preparation and Sequencing

Extracted DNA (2 ng DNA in 50 uL buffer) was sheared to 450bp using a Covaris LE220 system. Library preparation was performed using a Biomek FXP Automated Liquid Handling Workstation (Beckman Coulter) with the KAPA HyperPrep Kit (Roche), with 12 cycles of PCR. Final libraries were normalized and pooled, with 20 samples per poor. Each pool was run on 2 lanes of an Illumina HiSeq 4000 using the paired-end 2x150bp protocol. Library preparation and sequencing was performed at the New York University Langone Medical Center Genome Technology Core. Sequences can be found under the BioProject accession number PRJNA573963 (http://www.ncbi.nlm.nih.gov/bioproject/573963) for the levofloxacin dataset and under the BioProject accession number PRJNA531921 (https://www.ncbi.nlm.nih.gov/bioproject/531921) for the test dataset.

#### Data Processing

Raw shotgun sequencing reads were processed using Kneaddata (v0.6.1) to remove contaminating human sequences from the dataset[159]. Briefly, the *kneaddata* function was used with the Bowtie2 *Homo sapiens* database (v0.1)[160] to remove contaminating host reads from the sequencing files. See Additional File 1 for more information.

#### Taxonomic Classification

Kraken2, a taxonomic classifier that maps shotgun sequencing k-mers to genomic databases, was used to assign taxonomy to kneaddata-processed shotgun sequencing reads[70]. Briefly, the *kraken2-build* function was used to create a custom database containing the “bacteria” and “archaea” from NCBI libraries, and the *kraken2* function was used to run the kneaddata-filtered shotgun sequencing reads against this database and assign taxonomy. While Kraken2 does not estimate species abundances, Bracken2 (Bayesian Reestimation of Abundance with KrakEN) uses the taxonomy assigned by Kraken2 to estimate the number of reads per sample that originate from individual species[71]. The Kraken2 database was used to create a Bracken-compatible database using the *bracken-build* function, and the Kraken2 report files for each sample were run against the Bracken database using the *bracken* function for the phylum, genus, and species levels. Phylum- and species-level relative abundance outputs were formatted for biomarker discovery using LEfSe. The *kraken-biom* function was used to convert the Bracken report files into a biom file for import into R. Output data can be found in Additional Data 1. Relative abundance plots were generated in GraphPad Prism v8.

#### Taxonomic Diversity Analysis

Alpha and beta diversity analyses were performed using the phyloseq (v1.27.2)[161, 162] and vegan (v2.5-4)[163] packages in R (v3.4.3). Briefly, the biom file was imported into a phyloseq object. The phyloseq *estimate_richness* function was used to obtain Shannon’s Diversity Index values for all samples, while the vegan *phyloseq::distance* and *ordinate* functions were used to generate a Bray-Curtis matrix and PCoA values. See Additional File 1 for more information. Data was exported as csv files for formatting, and plotting was performed in GraphPad Prism v8.

#### Gene and Pathways Analysis

The Human Microbiome Project Unified Metabolic Analysis Network 2 (HUMAnN2) pipeline was used to profile the presence and abundance of genetic pathways in our samples[97]. Briefly, the *humann2* function was used with the kneaddata-filtered metagenomic sequences to estimate genes and MetaCyc pathways present in the samples based on the UniRef90 database, files were joined using the *humann2_join_tables* function and the full tables were de-leveled using the *humann2_split_stratified_table* function. The unstratified gene-level abundances were converted to both GO terms and KEGG orthologs using the *humann2_regroup_table* function, and the *humann2_renorm_table* function was used to normalize the MetaCyc pathway, GO term, and KEGG ortholog tables by computing relative abundance. These relative abundance tables were formatted for biomarker discovery with LEfSe. Additionally, the, and LEfSe was also used to analyze pre- and post-treated samples using both outputs. See Additional File 1 for more information. Output data can be found in Additional Data 3. Plots were generated in Graphpad Prism 8.

#### Resistome Analysis

The ARG content of the samples was analyzed using DeepARG-SS, a deep learning model that can predict ARGs from short-read metagenomic data[98]. We first analyzed the data using the *deeparg* function with the *-reads* flag. The mapped ARGs output was then imported into R, where it was processed to obtain tables of the ARGs detected per sample at both the specific gene and antibiotic class levels. The ARGs detected were normalized to the number of reads per sample.

Additionally, after metagenomic assembly and binning was performed (see below), individual bins were analyzed using DeepARG-LS, a deep learning model optimized to predict ARGS from gene-level input. The *DNA_features* output from selected bins was analyzed using the *deeparg* function with the *-genes* flag to analyze whether the levels or identity of ARGs could be linked to specific species of interest. The ARGs detected were normalized to the number of features per bin. All output data can be found in Additional Data 4. Plots were generated in GraphPad Prism 8.

#### Metagenomic Assembly and Binning

To further examine the ARGs present in the samples, kneaddata-filtered reads were uploaded to the web-based Pathosystems Resource Integration Center (PATRIC)[164]. Reads were assembled into contigs using the *auto* option of the Assembly service, which provides both raw output contigs from specific assembly algorithms and contigs of the “best” assembly as judged by the in-house PATRIC script ARAST. We ran the assembly using two different inputs: reads that had been processed by *kneaddata* as pairs, which has the advantage of utilizing mate-pairing information for longer total reads, and reads that had been processed by *kneaddata* after pairs were concatenated into a single file, which has the advantage of keeping reads whose mates failed trimming.

Both the raw SPAdes assembly algorithm contigs[165] and the best assembly contigs were then processed using the Metagenomics Binning service, which assigns contigs to specific organisms and annotates the bin’s genome. Quality measures were used to define bins as either “good”, “acceptable”, or “bad” according to the criteria in Additional Table 3, and only “good” or “acceptable” bins were used moving forward. When more than one binning strategy (paired assembly or single assembly, SPAdes contigs or best contigs) called a particular bin as “good” or “acceptable”, quality measures from the four strategies were compared and the highest-quality bin for a given species of interest was chosen for ARG analysis. Finally, only bins of species present at 0.1% or greater relative abundance in the corresponding sample were selected for further analysis. A list of bins used, their source, and quality measures can be found in Additional Table 4.

#### Taxonomic Biomarker Analysis

LEfSe was used to identify potential biomarkers distinguishing levofloxacin-treated samples[95]. In all cases (taxonomic abundances, MetaCyc pathways, KEGG orthologs, GO terms, ARGs), data was formatted into csv files and uploaded to the Galaxy webserver. LEfSe was run under default parameters for biomarker detection, comparing either all pre-levofloxacin to all post-levofloxacin or immediately pre-levofloxacin to immediately post-levofloxacin. LEfSe was also used to compare genus-level taxonomic abundance outputs from Kraken2/Bracken2 and QIIME2, again under default parameters.

## DECLARATIONS

### Ethics Approval and Consent to Participate

Written information about SPREAD was mailed to the healthcare proxies of all eligible subjects. Proxies were then telephoned two weeks later to solicit participation and verbally obtain informed consent for the participation of themselves and the subjects. Approval for SPREAD, including the consent procedures, was obtained from the Institutional Review Board committee at Hebrew SeniorLife.

### Consent for Publication

Not applicable

### Availability of Data and Material

The underlying sequencing data for the current study are available in the NCBI Short Read Archive repository; 16S rRNA and shotgun metagenomics data for the levofloxacin samples can be found at BioProject accession number PRJNA573963 (http://www.ncbi.nlm.nih.gov/bioproject/573963) and shotgun metagenomics data for the test dataset can be found at BioProject accession number PRJNA531921 (https://www.ncbi.nlm.nih.gov/bioproject/531921). Analysis code and data generated from this study can be found in Additional File 1 and Additional Data 1-4 of this published article, respectively.

### Author’s Contributions

ADR conceptualized the project, performed analysis, generated figures, and wrote the manuscript. RA collected the data and contributed to manuscript preparation. EMCD collected the data and contributed to manuscript preparation. PB conceptualized the project and wrote the manuscript. All authors read and approved the final manuscript.

## Supporting information

All Additional Figures

All Additional Tables

## Acknowledgements

The authors acknowledge the laboratory of Dr. Martin Blaser and the New York University Langone Medical Center Genome Technology Core, which prepared metagenomic libraries and performed all 16S rRNA and shotgun metagenomics sequencing for this study.

## ADDITIONAL FILES

*Additional Figure 1: Relative Abundances of Phyla Across and Within Subjects*

(A) Relative abundance of phyla in all samples, ranked by average across all samples. (B-L) Relative abundances of phyla by subject, ranked by average within each subject.

*Additional Figure 2: Relative Abundances of Species Across and Within Subjects*

(A) Relative abundance of species in all samples, grouped by genus and phylum and ranked within those levels by average relative abundance across all samples. (B-L) Relative abundances of phyla by subject, grouped by genus and phylum ranked within those levels by average within each subject. Coloring is the same as in Figure 2A.

*Additional Figure 3: Comparison of Genus-level Classifications by Metagenomics and 16S rRNA Analysis*

(A) Relative abundances of genera called by both QIIME2 and Kraken2/Bracken2, where pairs of stacked bars indicate the same sample as measured by both methods. (B) Genera called by LEfSe as associated with either QIIME2 or Kraken2/Bracken2. Each genus is colored according to its source phylum.

*Additional Figure 4: Relative Abundance of Gene Ontology Terms Across All Samples*

(A) Relative abundances of the top 250 most-abundant GO terms, representing broad functional categories, across all samples. A significant proportion are “unmapped” or “ungrouped”, as not all UniRef90 gene families can be mapped to a GO term.

*Additional Figure 5: Relative Abundance of Antimicrobial Resistance Genes Within and Across Subjects*

(A) Relative abundance of species in all samples, grouped and ranked within class by average relative abundance across all samples. (B-L) Relative abundances of ARGs by subject, grouped and ranked within class by average relative abundance within each subject. Coloring is the same as in Figure 2B.

*Additional Figure 6: Comparison of MDRO and non-MDRO Bins of the Same Species*

(A) ARG density in all E. coli bins across samples. (B) ARG density in all P. mirabilis bins across samples. (C) Beta-lactam ARG density in all S. aureus bins across samples.

*Additional Table 1: Metadata on levofloxacin cohort from SPREAD*

This table lists the age, biological sex, and race of all subjects, whether a multidrug-resistant organism (MDRO) was detected in the subject at any timepoint, the duration of levofloxacin administration, and the reason for which they were administered levofloxacin. For MDROs, the specific organism detected and the antimicrobial agents it was found to be resistant to are also listed.

*Additional Table 2: Overview of longitudinal sample collection from levofloxacin cohort from SPREAD*

This table lists all samples from the levofloxacin cohort that were collected, sequenced, or analyzed in this study. Samples that were successfully analyzed are marked with a “yes”, while samples that could not be collected, sequenced, or analyzed are marked with a “no”. For samples that were not analyzed, a reason is also provided according to the following key: SD = <u>s</u>ubject <u>d</u>eceased at this timepoint, NC = sample was <u>n</u>ot <u>c</u>ollected, NS = sample was <u>n</u>ot <u>s</u>equenced, SP = sample <u>s</u>equenced <u>p</u>oorly.

*Additional Table 3: Bin selection quality cutoffs*

This table lists the cutoffs used to determine whether a bin was “good” or “acceptable” to be used in further analysis, or “bad” enough to be discarded. Briefly, “good” bins had to meet the “good” cutoffs for all five criteria measured, “acceptable” bins could have a maximum of two “acceptable” criteria as long as all others were “good”, and “bad” bins contained any criterion below the “bad” cutoffs.

*Additional Table 4: Bins selected for DeepARG analysis*

This table lists all of the bins generated by PATRIC that were selected based on the criteria in Additional Table 3 to be analyzed using DeepARG. It includes all quality scores used to assess bin quality, as well as the PATRIC reference genome used to annotate the bin.

*Additional Table 5: BioProject Sample Identifiers for Test Dataset*

This table lists the sample names used in this study, the SPREAD IDs, and the BioProject PRJNA531921 sample names for the shotgun metagenomics sequencing files of the 67-sample dataset used to test the multiple linear regression developed from the levofloxacin dataset.

*Additional Data 1: Taxonomic Classifications from Shotgun Metagenomics Kraken2/Bracken2*

This file includes the relative abundances of the taxonomic classifications at the phylum, genus, and species level for both the initial levofloxacin-treated dataset (tabs 1, 2, and 3) and the second, larger test dataset (tabs 4, 5, and 6). Additional Table 5 links the sample names used for the test dataset in this study with their identifiers in BioProject PRJNA531921.

*Additional Data 2: Taxonomic Classifications from 16S rRNA Sequencing QIIME2*

This file includes the relative abundances of the taxonomic classifications at the phylum (tab 1) and genus (tab 2) level for the initial levofloxacin-treated dataset.

*Additional Data 3: Metagenomic Classifications from HUMANn2*

This file includes the relative abundances of the MetaCyc pathway (tab 1), KEGG ortholog (tab 2), and GO term (tab 3) outputs for the initial levofloxacin-treated dataset.

*Additional Data 4: Antimicrobial Resistance Gene profiles from DeepARG*

This file includes the relative abundances of antimicrobial class and specific resistance genes at the phylum, genus, and species level for the initial levofloxacin-treated dataset (tabs 1 and 2), the second, larger test dataset (tabs 3 and 4), and the bins generated by PATRIC (tabs 5 and 6). Additional Table 5 links the sample names used for the test dataset in this study with their identifiers in BioProject PRJNA531921.

*Additional File 1: Code Used for Analysis*

This file includes all of the analysis code used for QIIME2, Kraken2 and Bracken, Phyloseq, HUMANn2, and DeepARG.

