## Supplementary material for "Antimicrobial resistance gene prevalence in a population of patients with advanced dementia is related to specific pathobionts": All Additional Figures

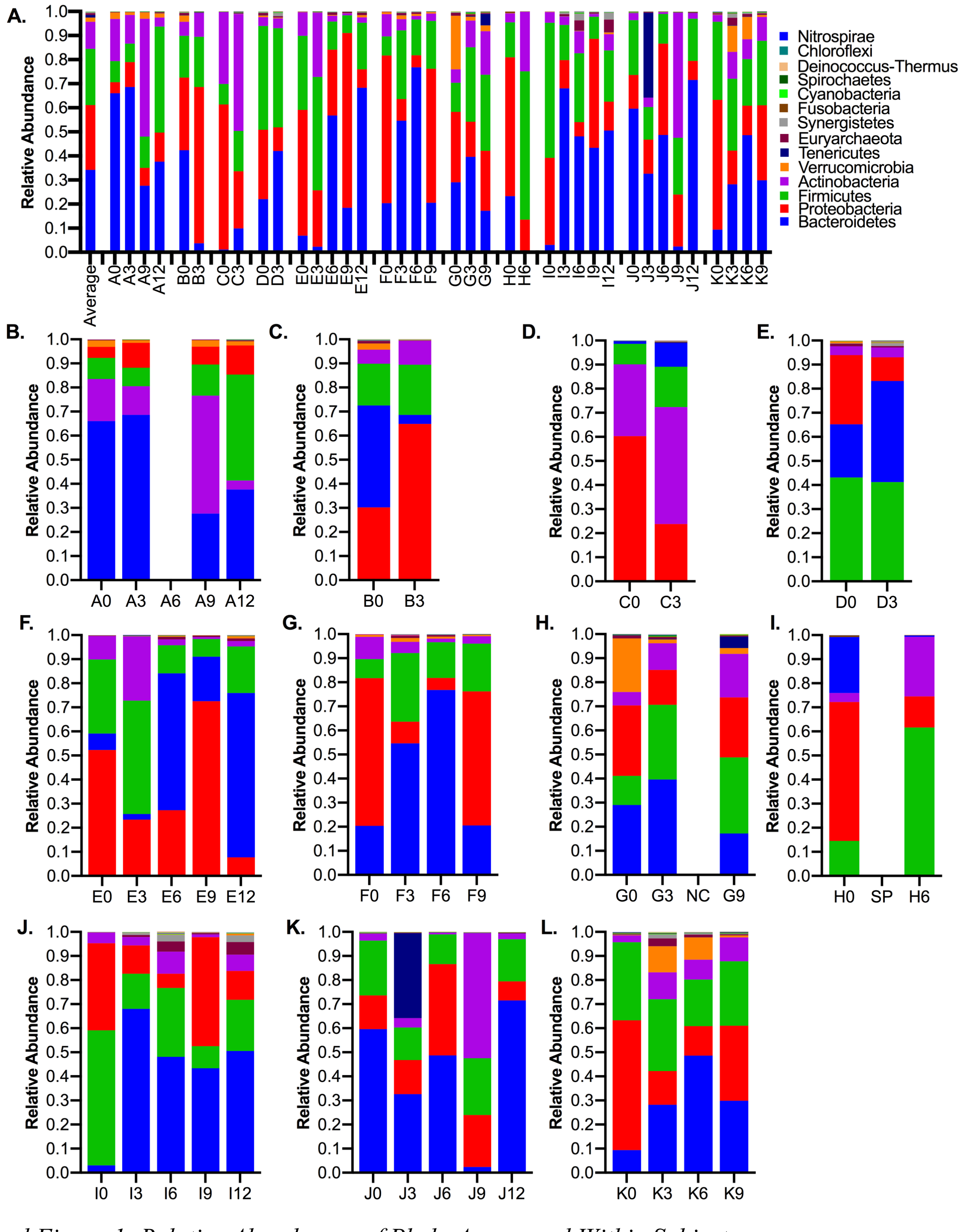


*Additional Figure 1: Relative Abundances of Phyla Across and Within Subjects*

(A) Relative abundance of phyla in all samples, ranked by average across all samples. (B-L) Relative abundances of phyla by subject, ranked by average within each subject.

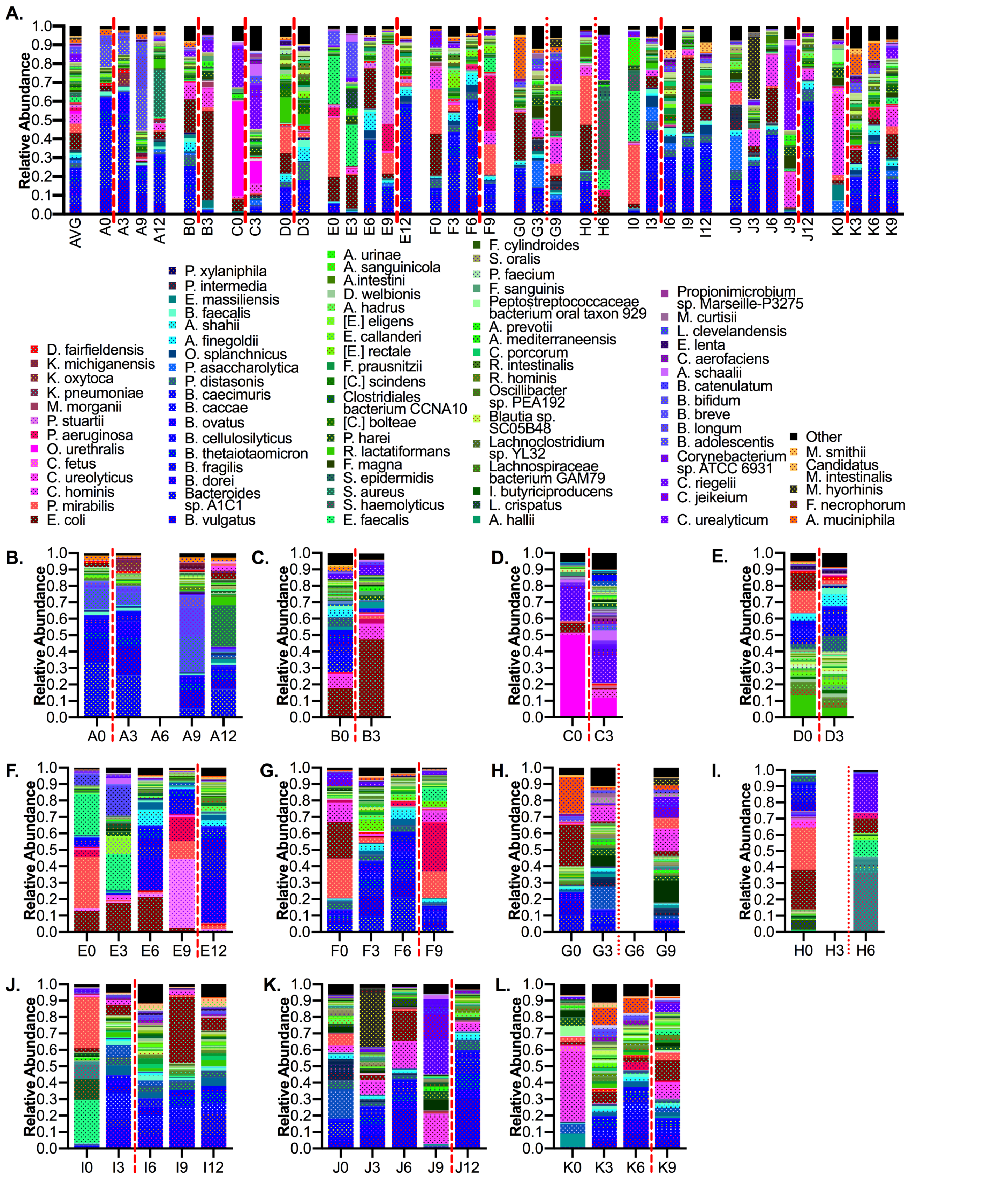


*Additional Figure 2: Relative Abundances of Species Across and Within Subjects*

(A) Relative abundance of species in all samples, grouped by genus and phylum and ranked within those levels by average relative abundance across all samples. (B-L) Relative abundances of phyla by subject, grouped by genus and phylum ranked within those levels by average within each subject. Coloring is the same as in Figure 2A.

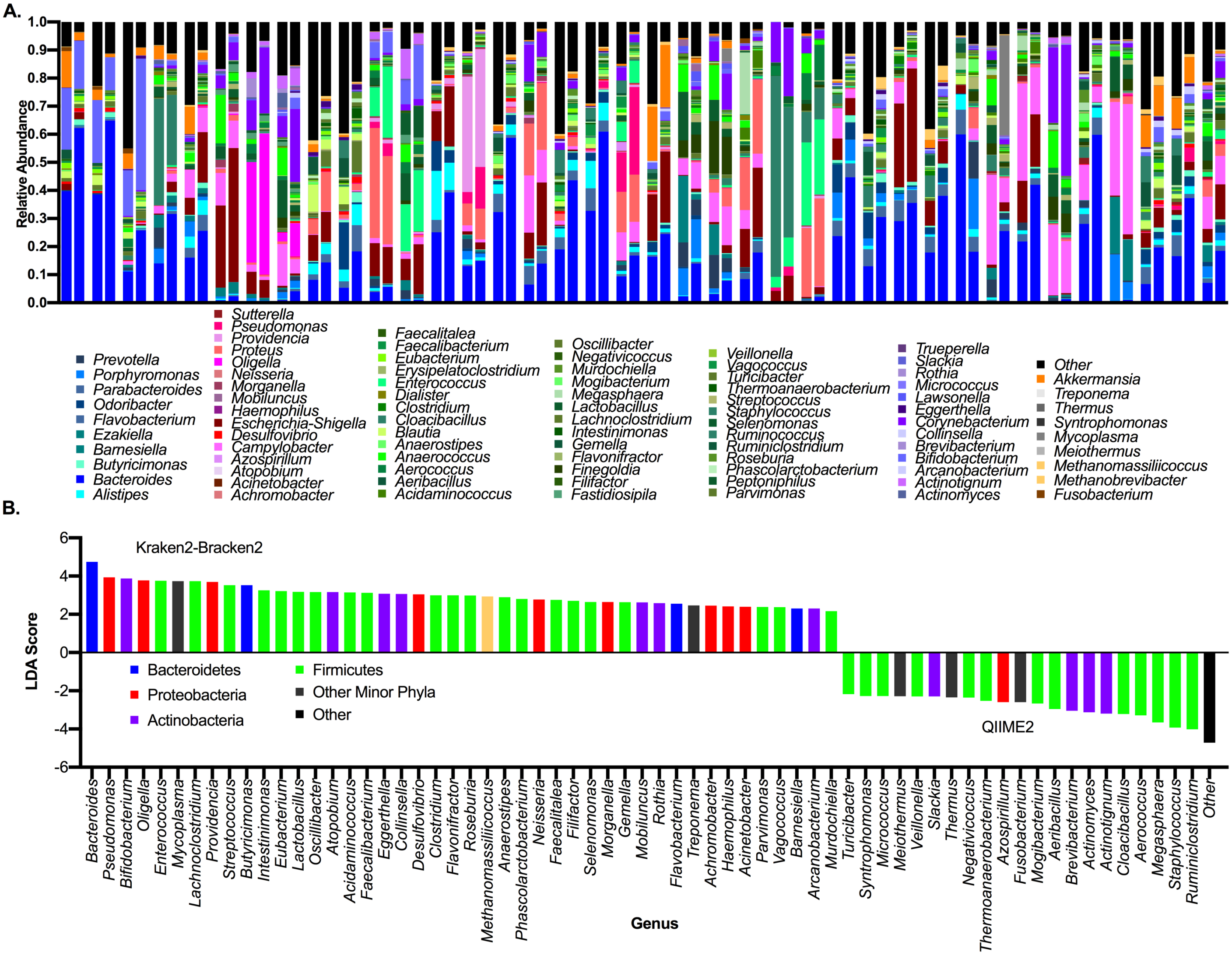


*Additional Figure 3: Comparison of Genus-level Classifications by Metagenomics and 16S rRNA Analysis*
(A) Relative abundances of genera called by both QIIME2 and Kraken2/Bracken2, where pairs of stacked bars indicate the same sample as measured by both methods. (B) Genera called by LEfSe as associated with either QIIME2 or Kraken2/Bracken2. Each genus is colored according to its source phylum.

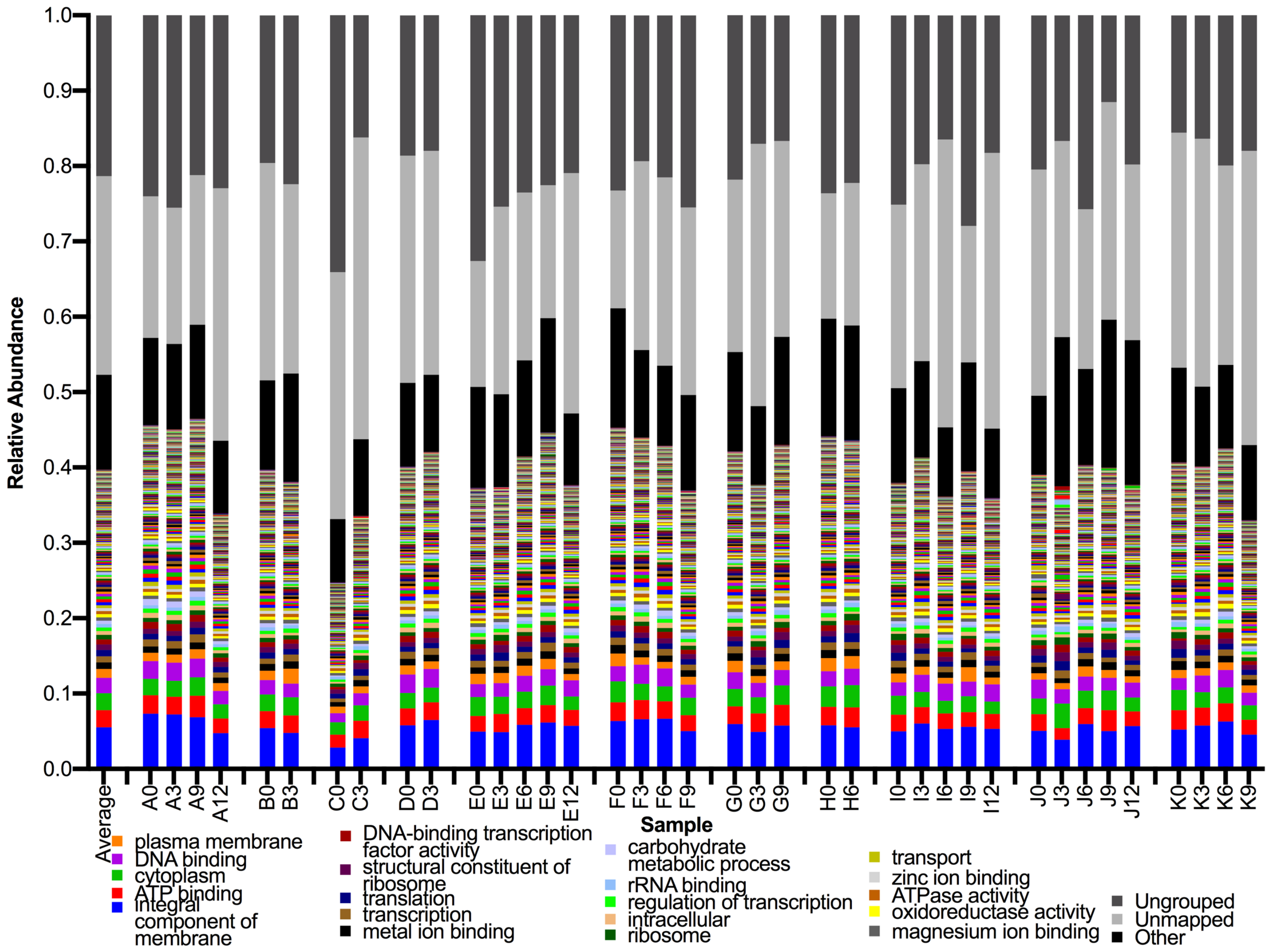


*Additional Figure 4: Relative Abundance of Gene Ontology Terms Across All Samples*

(A) Relative abundances of the top 250 most-abundant GO terms, representing broad functional categories, across all samples. A significant proportion are “unmapped” or “ungrouped”, as not all UniRef90 gene families can be mapped to a GO term.

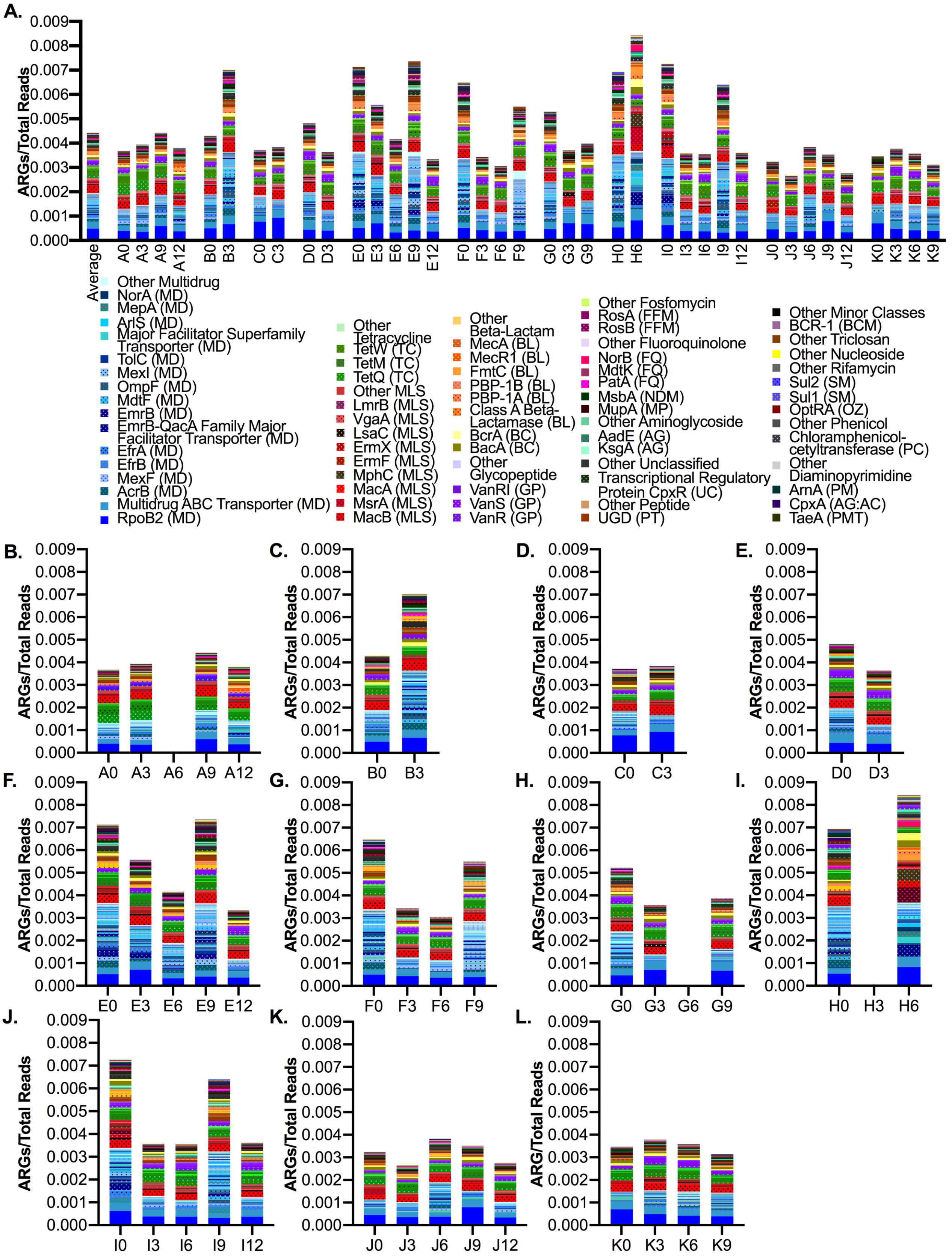


*Additional Figure 5: Relative Abundance of Antimicrobial Resistance Genes Within and Across Subjects*
(A) Relative abundance of species in all samples, grouped and ranked within class by average relative abundance across all samples. (B-L) Relative abundances of ARGs by subject, grouped and ranked within class by average relative abundance within each subject. Coloring is the same as in Figure 2B.

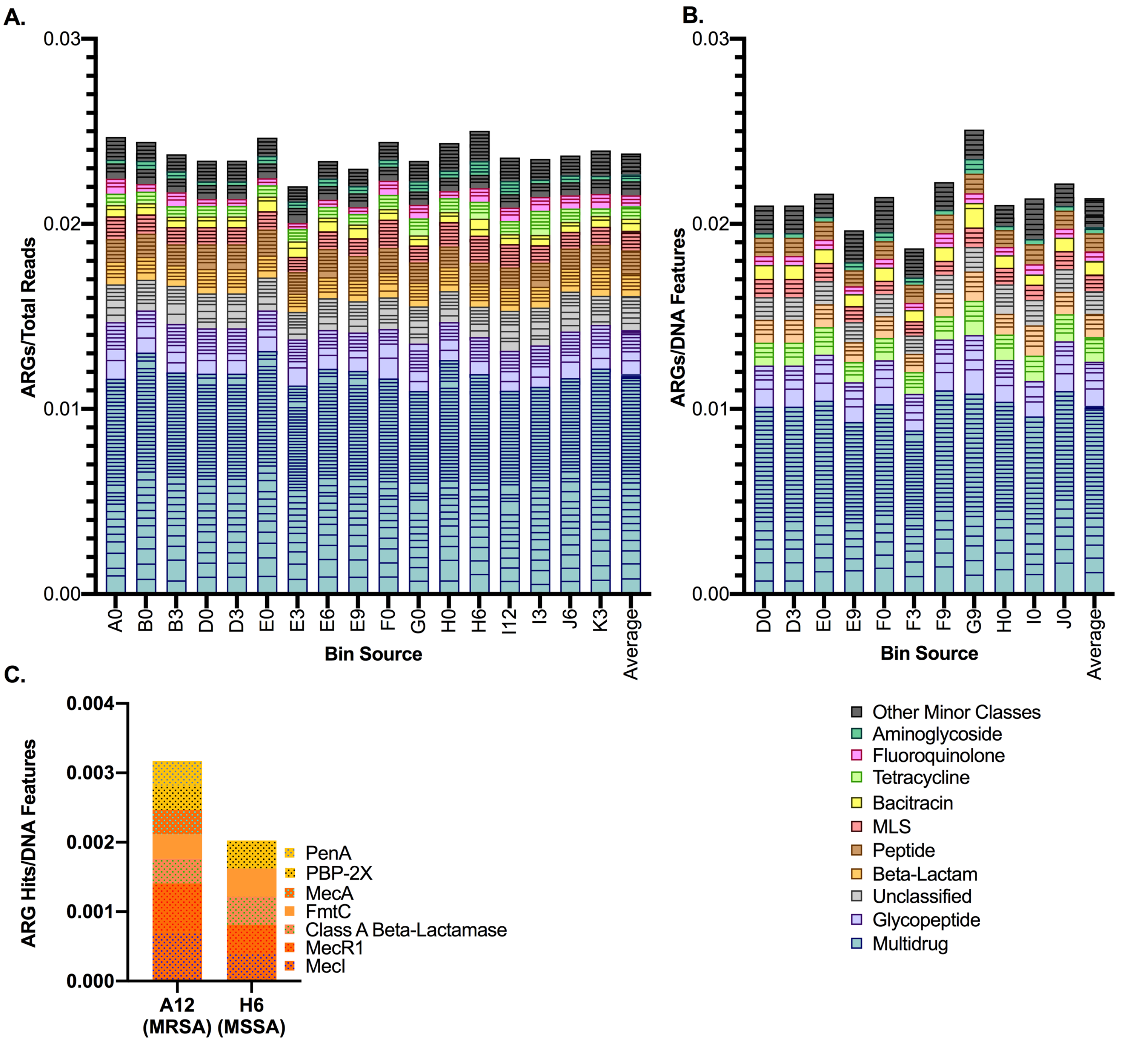


*Additional Figure 6: Comparison of MDRO and non-MDRO Bins of the Same Species*

(A) ARG density in all *E. coli* bins across samples. (B) ARG density in all *P. mirabilis* bins across samples. (C) Beta-lactam ARG density in all *S. aureus* bins across samples.
