## Supplementary material for "Antimicrobial resistance gene prevalence in a population of patients with advanced dementia is related to specific pathobionts": All Additional Tables

| Subject | Sex | Age | Race | MDRO Detected | Levofloxacin Duration | Reason for Levofloxacin Administration |
|---|---|---|---|---|---|---|
| **A** | F | 94 | W | Yes (*S. aureus*; methicillin) | 7 days | Urinary tract infection |
| **B** | F | 101 | W | Yes (*E. coli*; ampicillin/sulbactam, cefazolin, ceftazidime, ceftriaxone, ciprofloxacin) | 6 days | Upper respiratory tract infection |
| **C** | F | 88 | W | Yes (*P. mirabilis*; ampicillin/sulbactam, ciprofloxacin, gentamicin) | 7 days | Urinary tract infection |
| **D** | F | 74 | W | Yes (*P. mirabilis*; ampicillin/sulbactam, ciprofloxacin, gentamicin) | 10 days | Upper respiratory tract infection |
| **E** | F | 78 | W | No | 7 days | Upper respiratory tract infection |
| **F** | F | 101 | W | No | 10 days | Upper respiratory tract infection |
| **G** | F | 83 | NW | No | 11 days | Fever of unknown source |
| **H** | F | 87 | W | No | 10 days | Upper respiratory tract infection |
| **I** | M | 89 | W | No | 8 days | Upper respiratory tract infection |
| **J** | F | 86 | W | No | 7 days | Fever of unknown source |
| **K** | F | 91 | W | No | 6 days | Upper respiratory tract infection |

*Additional Table 1: Metadata on levofloxacin cohort from SPREAD*

This table lists the age, biological sex, and race of all subjects, whether a multidrug-resistant organism (MDRO) was detected in the subject at any timepoint, the duration of levofloxacin administration, and the reason for which they were administered levofloxacin. For MDROs, the specific organism detected and the antimicrobial agents it was found to be resistant to are also listed.

| Subject | T0 | T3 | T6 | T9 | T12 |
|---|---|---|---|---|---|
| **A** | Yes | Yes | No – NS | Yes | Yes |
| **B** | Yes | Yes | No – SD | No – SD | No – SD |
| **C** | Yes | Yes | No – SD | No – SD | No – SD |
| **D** | Yes | Yes | No - SD | No - SD | No - SD |
| **E** | Yes | Yes | Yes | Yes | Yes |
| **F** | Yes | Yes | Yes | Yes | No - NS |
| **G** | Yes | Yes | No - NC | Yes | No – SD |
| **H** | Yes | No - SP | Yes | No - SD | No – SD |
| **I** | Yes | Yes | Yes | Yes | Yes |
| **J** | Yes | Yes | Yes | Yes | Yes |
| **K** | Yes | Yes | Yes | Yes | No - SD |

*Additional Table 2: Overview of longitudinal sample collection from levofloxacin cohort from SPREAD*

This table lists all samples from the levofloxacin cohort that were collected, sequenced, or analyzed in this study. Samples that were successfully analyzed are marked with a "yes", while samples that could not be collected, sequenced, or analyzed are marked with a "no". For samples that were not analyzed, a reason is also provided according to the following key: SD = subject deceased at this timepoint, NC = sample was not collected, NS = sample was not sequenced, SP = sample sequenced poorly.

| **Bin Type** | **Coarse Consistency** | **Completeness** | **Fine Consistency** | **Contamination** | **Single PheS** | **Number of Criteria to be Met** |
|---|---|---|---|---|---|---|
| Good | 87%+ | 87%+ | 87%+ | <10% | Yes | All 5 |
| Acceptable | 80-86.9% | 80-86.9% | 80-64.9% | 10.1-20% | No | 1-2, if all others are "good" |
| Bad | <80% | <80% | <80% | 20%+ | No | Any "bad" criterion or 3+ "acceptable" criteria |

*Additional Table 3: Bin selection quality cutoffs*

This table lists the cutoffs used to determine whether a bin was "good" or "acceptable" to be used in further analysis, or "bad" enough to be discarded. Briefly, "good" bins had to meet the "good" cutoffs for all five criteria measured, "acceptable" bins could have a maximum of two "acceptable" criteria as long as all others were "good", and "bad" bins contained any criterion below the "bad" cutoffs.

| Read Type | Binning Strategy | Source Sample | PATRIC Score | Species Identification | Reference Genome | FC | CP | CC | CM | PheS |
|---|---|---|---|---|---|---|---|---|---|---|
| Single | Best | A0 | 1528 | *Bacteroides ovatus* | 28116.191 | 98.2 | 96.4 | 92.3 | 8.9 | Yes |
| Paired | All | A0 | 1257 | *Bacteroides thetaiotaomicron* | 818.2 | 96 | 93.7 | 82.1 | 11.7 | Yes |
| Single | Best | A0 | 1829 | *Bacteroides vulgatus* | 821.509 | 98.5 | 97.5 | 96.2 | 3.9 | Yes |
| Single | All | A0 | 1978 | *Bifidobacterium adolescentis* | 1680.104 | 99.3 | 98.4 | 98.6 | 1.6 | Yes |
| Single | Best | A0 | 1766 | *Bifidobacterium bifidum* | 500634.3 | 97.9 | 96.6 | 94.9 | 4.7 | Yes |
| Single | Best | A0 | 1648 | *Bifidobacterium longum* | 1298922.3 | 98.8 | 96.6 | 98.6 | 7.8 | Yes |
| Single | Best | A0 | 2017 | *Escherichia coli* | 562.30402 | 99.5 | 98.4 | 100 | 1.1 | Yes |
| Single | Best | A0 | 1583 | *Eubacterium callanderi* | 53442.4 | 99 | 94.4 | 100 | 8.9 | Yes |
| Single | Best | A3 | 1711 | *Bacteroides ovatus* | 28116.191 | 99 | 97.3 | 98.1 | 6.6 | Yes |
| Paired | Best | A3 | 1311 | *Bacteroides thetaiotaomicron* | 818.2 | 96.5 | 94.8 | 80.8 | 10.6 | Yes |
| Single | Best | A3 | 1789 | *Bacteroides vulgatus* | 821.509 | 98.5 | 97.7 | 99 | 5.3 | Yes |
| Single | Best | A3 | 1935 | *Bifidobacterium adolescentis* | 1680.104 | 99 | 98.2 | 98.5 | 2.4 | Yes |
| Paired | Best | A3 | 1729 | *Bifidobacterium longum* | 1298922.3 | 99.6 | 96.7 | 100 | 6.5 | Yes |
| Single | Best | A3 | 1844 | *Klebsiella oxytoca* | 571.142 | 99.6 | 94.9 | 100 | 3.8 | Yes |
| Single | All | A9 | 1119 | *Bacteroides ovatus* | 28116.191 | 95.6 | 89.6 | 85.3 | 14.2 | Yes |
| Single | Best | A9 | 1590 | *Bacteroides thetaiotaomicron* | 818.2 | 98.2 | 95.2 | 92.3 | 7.4 | Yes |
| Single | Best | A9 | 1678 | *Bacteroides vulgatus* | 821.509 | 98.7 | 97.1 | 99.5 | 7.5 | Yes |
| Single | Best | A9 | 2029 | *Bifidobacterium adolescentis* | 1680.104 | 99.6 | 99 | 100 | 1 | Yes |
| Single | Best | A9 | 1985 | *Bifidobacterium bifidum* | 500634.3 | 99.3 | 98.9 | 99.7 | 1.8 | Yes |
| Paired | Best | A9 | 1672 | *Bifidobacterium longum* | 1298922.3 | 99.3 | 97 | 100 | 7.7 | Yes |
| Single | Best | A12 | 2055 | *Staphylococcus aureus* | 1280.1372 | 100 | 99.6 | 100 | 0.6 | Yes |

| | | | | | | | | | | |
|---|---|---|---|---|---|---|---|---|---|---|
| Single | Best | B0 | 1264 | *Corynebacterium urealyticum* | 43771.9 | 96.7 | 92.4 | 99.2 | 14.7 | Yes |
| Single | Best | B0 | 2039 | *Escherichia coli* | 562.23379 | 99.9 | 98.6 | 100 | 0.7 | Yes |
| Single | Best | B0 | 2012 | *Parabacteroides distasonis* | 823.7 | 99.3 | 97.5 | 100 | 1 | Yes |
| Single | All | B0 | 1806 | *Methanobrevibacter smithii* | 1263088.3 | 99.4 | 98 | 98.3 | 4.9 | Yes |
| Single | All | B0 | 1455 | *Porphyromonas asaccharolytica* | 879243.3 | 96.5 | 93.4 | 98.7 | 11 | Yes |
| Paired | Best | B0 | 1199 | *Alistipes finegoldii* | 214856.4 | 98.4 | 92.6 | 99.5 | 16.1 | Yes |
| Single | Best | B3 | 1416 | *Corynebacterium urealyticum* | 43771.9 | 94.3 | 92.1 | 87.8 | 9.3 | Yes |
| Paired | Best | B3 | 1877 | *Escherichia coli* | 562.17562 | 99.7 | 97 | 100 | 3.6 | No |
| Paired | Best | B3 | 1770 | *Fastidiosipila sanguinis* | 236753.3 | 93.3 | 92.8 | 86.9 | 2.2 | Yes |
| Single | Best | C3 | 1488 | *Corynebacterium urealyticum* | 43771.9 | 95.1 | 92.2 | 95.4 | 9.4 | Yes |
| Single | Best | D0 | 1252 | *[Eubacterium] eligens* | 39485.21 | 99 | 94.3 | 100 | 15.5 | Yes |
| Single | Best | D0 | 1891 | *Bacteroides fragilis* | 817.199 | 96.6 | 95.6 | 93.9 | 1.8 | Yes |
| Single | Best | D0 | 1357 | *Bacteroides ovatus* | 28116.19 | 99.3 | 94.5 | 98.7 | 13.2 | Yes |
| Single | Best | D0 | 1108 | *Phascolarctobacterium faecium* | 1122957.3 | 95.6 | 91.4 | 98.7 | 17.5 | Yes |
| Single | Best | D0 | 2081 | *Methanobrevibacter smithii* | 420247.28 | 100 | 99.7 | 100 | 0.1 | Yes |
| Paired | Best | D0 | 1919 | *Eggerthella lenta* | 84112.24 | 98.3 | 95.3 | 96.6 | 1.7 | Yes |
| Paired | Best | D0 | 2072 | *Escherichia coli* | 562.28676 | 99.6 | 98.4 | 100 | 0 | Yes |
| Paired | Best | D0 | 1604 | *Parabacteroides distasonis* | 823.7 | 97 | 93.3 | 92.3 | 6.7 | Yes |
| Paired | Best | D0 | 2012 | *Proteus mirabilis* | 584.9 | 98.7 | 97 | 100 | 0.9 | Yes |
| Paired | Best | D3 | 1876 | *Bacteroides fragilis* | 817.199 | 96.7 | 95.2 | 93.9 | 2 | Yes |
| Paired | Best | D3 | 1078 | *Bacteroides ovatus* | 28116.19 | 98.7 | 94.4 | 94 | 17.8 | Yes |
| Paired | Best | D3 | 1999 | *Methanobrevibacter smithii* | 420247.28 | 99.9 | 99.5 | 100 | 1.7 | Yes |
| Paired | Best | D3 | 1604 | *Parabacteroides distasonis* | 823.7 | 97 | 93.3 | 92.3 | 6.7 | Yes |

| | | | | | | | | | | |
|---|---|---|---|---|---|---|---|---|---|---|
| Paired | Best | D3 | 2012 | *Proteus mirabilis* | 584.9 | 98.7 | 97 | 100 | 0.9 | Yes |
| Paired | Best | D3 | 2072 | *Escherichia coli* | 562.28676 | 99.6 | 98.4 | 100 | 0 | Yes |
| Paired | Best | D3 | 1919 | *Eggerthella lenta* | 84112.24 | 98.3 | 95.3 | 96.6 | 1.7 | Yes |
| Single | Best | D3 | 1116 | *Eubacterium rectale* | 1263079.3 | 98.7 | 93.1 | 96.7 | 17.3 | Yes |
| Single | Best | D3 | 1293 | *Intestinimonas butyriciproducens* | 1297617.27 | 93.9 | 89.1 | 88.2 | 11.2 | Yes |
| Single | Best | E0 | 1802 | *Bacteroides dorei* | 997877.5 | 99.8 | 97.9 | 100 | 5.3 | Yes |
| Single | Best | E0 | 2079 | *Bifidobacterium breve* | 1385939.3 | 99.5 | 99 | 100 | 0 | Yes |
| Single | Best | E0 | 2012 | *Escherichia coli* | 562.22574 | 99.7 | 97.9 | 100 | 1.1 | Yes |
| Single | Best | E0 | 1972 | *Mobiluncus curtisii* | 887899.3 | 96.3 | 95.6 | 100 | 1.4 | Yes |
| Single | Best | E0 | 2017 | *Peptoniphilus harei* | 54005.3 | 97.6 | 96.6 | 100 | 0.7 | Yes |
| Single | Best | E0 | 2012 | *Proteus mirabilis* | 584.293 | 98.8 | 97 | 100 | 0.9 | Yes |
| Single | Best | E0 | 2047 | *Pseudomonas aeruginosa* | 1402503.3 | 99.6 | 98 | 99.4 | 0.3 | Yes |
| Paired | Best | E0 | 1880 | *Enterococcus faecalis* | 1351.868 | 99.6 | 97.3 | 100 | 3.6 | Yes |
| Single | Best | E3 | 2065 | *Bifidobacterium breve* | 1385939.3 | 99.6 | 99.1 | 100 | 0.3 | Yes |
| Single | Best | E3 | 1339 | *Escherichia coli* | 562.22574 | 98.4 | 87.6 | 100 | 12.3 | Yes |
| Single | Best | E3 | 1669 | *Fastidiosipila sanguinis* | 236753.3 | 92.9 | 92.5 | 87.6 | 4.3 | Yes |
| Single | Best | E3 | 2072 | *Lactobacillus crispatus* | 47770.179 | 99.7 | 98.4 | 100 | 0 | Yes |
| Single | Best | E3 | 1131 | *Peptoniphilus harei* | 54005.3 | 97.5 | 92.3 | 100 | 17.5 | Yes |
| Single | Best | E6 | 1292 | *Anaerostipes hadrus* | 649756.24 | 97 | 92.2 | 99.3 | 14.1 | Yes |
| Single | Best | E6 | 2034 | *Bacteroides dorei* | 997877.5 | 98.7 | 98.4 | 96.2 | 0 | Yes |
| Single | Best | E6 | 2088 | *Methanobrevibacter smithii* | 420247.28 | 99.9 | 99.9 | 100 | 0 | Yes |
| Single | Best | E6 | 1912 | *Providencia stuartii* | 588.6 | 98.7 | 94.2 | 98.6 | 2 | Yes |
| Paired | Best | E6 | 2060 | *Bacteroides fragilis* | 817.25 | 98.8 | 97.6 | 99.7 | 0 | Yes |
| Paired | Best | E6 | 1732 | *Bacteroides thetaiotaomicron* | 818.2 | 98.4 | 96.2 | 94.9 | 5.3 | Yes |
| Paired | Best | E6 | 2032 | *Escherichia coli* | 562.22574 | 99.8 | 97.9 | 100 | 0.7 | Yes |
| Paired | Best | E6 | 2069 | *Odoribacter splanchnicus* | 28118.6 | 99.1 | 98.5 | 99.6 | 0 | Yes |

| | | | | | | | | | | |
|---|---|---|---|---|---|---|---|---|---|---|
| Paired | Best | E9 | 2012 | *Acidaminococcus intestini* | 1120921.3 | 97.3 | 96.7 | 99.3 | 0.7 | Yes |
| Paired | Best | E9 | 2017 | *Escherichia coli* | 562.22574 | 99.7 | 97.5 | 100 | 0.9 | Yes |
| Paired | Best | E9 | 1866 | *Bacteroides dorei* | 997877.5 | 99.8 | 99 | 98.7 | 4 | Yes |
| Paired | Best | E9 | 1883 | *Bacteroides fragilis* | 817.25 | 99 | 97.6 | 100 | 3.6 | Yes |
| Paired | Best | E9 | 1755 | *Bacteroides thetaiotaomicron* | 818.2 | 98.4 | 97.2 | 95.1 | 5.1 | Yes |
| Paired | Best | E9 | 2010 | *Campylobacter hominis* | 360107.7 | 99.7 | 99.3 | 97.3 | 0.9 | Yes |
| Paired | Best | E9 | 1937 | *Finegoldia magna* | 1260.9 | 99.9 | 98.4 | 100 | 2.7 | Yes |
| Paired | Best | E9 | 2056 | *Proteus mirabilis* | 584.293 | 98.7 | 96.9 | 100 | 0 | Yes |
| Paired | Best | E9 | 1986 | *Providencia stuartii* | 588.6 | 99 | 96.5 | 100 | 1.3 | Yes |
| Paired | Best | E9 | 2057 | *Pseudomonas aeruginosa* | 287.2475 | 99.7 | 98.7 | 99.7 | 0.3 | Yes |
| Single | Best | E9 | 1797 | *Alistipes finegoldii* | 214856.4 | 98.9 | 97 | 100 | 5.2 | Yes |
| Single | Best | E9 | 1704 | *Campylobacter ureolyticus* | 883165.3 | 96.5 | 94.5 | 93.9 | 5.3 | Yes |
| Single | Best | E9 | 1800 | *Peptoniphilus harei* | 54005.3 | 97.7 | 95 | 100 | 4.7 | Yes |
| Single | All | E9 | 1376 | *Fastidiosipila sanguinis* | 236753.3 | 93.1 | 91.5 | 87.9 | 10 | Yes |
| Single | All | E9 | 2083 | *Methanobrevibacter smithii* | 420247.28 | 99.9 | 99.4 | 100 | 0 | Yes |
| Paired | Best | E12 | 1980 | *Odoribacter splanchnicus* | 28118.6 | 99.5 | 97.7 | 99.6 | 1.6 | Yes |
| Single | Best | E12 | 1758 | *Akkermansia muciniphila* | 239935.94 | 98.1 | 94.4 | 100 | 5.4 | Yes |
| Single | Best | E12 | 1581 | *Anaerostipes hadrus* | 649756.24 | 98.4 | 95.3 | 99.3 | 9 | Yes |
| Single | Best | E12 | 1957 | *Bacteroides dorei* | 997877.5 | 99.9 | 99.3 | 100 | 2.5 | Yes |
| Single | Best | E12 | 1701 | *Bacteroides fragilis* | 817.25 | 99.3 | 97.6 | 98.3 | 6.9 | Yes |
| Single | Best | E12 | 1765 | *Bacteroides thetaiotaomicron* | 818.2 | 98 | 95.9 | 93.5 | 4.3 | Yes |
| Single | All | F0 | 1380 | *Akkermansia muciniphila* | 239935.96 | 99.2 | 93.2 | 100 | 12.7 | No |
| Single | Best | F0 | 1618 | *Anaerostipes hadrus* | 649756.5 | 98.2 | 95.3 | 100 | 8.4 | Yes |
| Single | Best | F0 | 1967 | *Proteus mirabilis* | 584.299 | 98.8 | 97 | 100 | 1.8 | Yes |

| | | | | | | | | | | |
|---|---|---|---|---|---|---|---|---|---|---|
| Single | Best | F0 | 1751 | *Morganella morganii* | 582.171 | 93.6 | 91.3 | 87.1 | 2.3 | Yes |
| Paired | Best | F0 | 1906 | *Campylobacter hominis* | 360107.7 | 99.8 | 99.4 | 97.3 | 3 | Yes |
| Paired | Best | F0 | 1707 | *Corynebacterium jeikeium* | 38289.3 | 97 | 94.3 | 100 | 6.4 | Yes |
| Paired | Best | F0 | 1626 | *Escherichia coli* | 562.23379 | 99 | 93.3 | 100 | 7.8 | Yes |
| Paired | Best | F0 | 2079 | *Finegoldia magna* | 1260.9 | 99.4 | 99 | 100 | 0 | Yes |
| Paired | Best | F0 | 1950 | *Parabacteroides distasonis* | 823.7 | 99.3 | 96.7 | 99.1 | 1.9 | Yes |
| Paired | Best | F0 | 1296 | *Peptoniphilus harei* | 54005.7 | 97.3 | 94 | 99.7 | 14.5 | Yes |
| Paired | Best | F3 | 1474 | *[Eubacterium] eligens* | 39485.21 | 96.7 | 94.8 | 89.1 | 9 | Yes |
| Paired | Best | F3 | 1457 | *Anaerostipes hadrus* | 649756.24 | 93.1 | 90.6 | 81 | 6.8 | Yes |
| Paired | Best | F3 | 1845 | *Bacteroides cellulosilyticus* | 246787.5 | 97.7 | 95.3 | 92.2 | 2.3 | Yes |
| Paired | Best | F3 | 1754 | *Eubacterium rectale* | 657317.3 | 99.3 | 97.4 | 99.3 | 6 | Yes |
| Single | All | F3 | 1091 | *Bacteroides caccae* | 47678.6 | 95.6 | 88.2 | 83 | 14 | Yes |
| Single | All | F3 | 1292 | *Campylobacter hominis* | 360107.7 | 99.3 | 95.8 | 97.3 | 14.5 | Yes |
| Single | All | F3 | 1395 | *Campylobacter ureolyticus* | 883165.3 | 93.8 | 91.2 | 88.6 | 9.7 | Yes |
| Single | Best | F3 | 1763 | *Akkermansia muciniphila* | 239935.96 | 99.2 | 97.6 | 97.5 | 5.5 | Yes |
| Single | Best | F3 | 1310 | *Alistipes finegoldii* | 1263035.3 | 96.8 | 93.3 | 91.9 | 12.5 | Yes |
| Single | Best | F3 | 1345 | *Bacteroides fragilis* | 817.199 | 93.6 | 90.5 | 89.4 | 10.7 | Yes |
| Single | Best | F3 | 1195 | *Bacteroides thetaiotaomicron* | 818.287 | 98 | 94.5 | 90.5 | 14.8 | Yes |
| Single | Best | F3 | 1655 | *Corynebacterium urealyticum* | 43771.9 | 96.9 | 95 | 99.5 | 7.5 | Yes |
| Single | Best | F3 | 1872 | *Odoribacter splanchnicus* | 28118.4 | 99.6 | 97.9 | 100 | 3.9 | Yes |
| Single | Best | F3 | 2030 | *Parabacteroides distasonis* | 823.7 | 98.2 | 95 | 100 | 0.1 | Yes |
| Single | Best | F3 | 1998 | *Proteus mirabilis* | 584.299 | 99 | 97.1 | 100 | 1.2 | Yes |
| Single | Best | F6 | 994 | *Alistipes finegoldii* | 1263035.3 | 95.9 | 92.4 | 89.7 | 18.2 | Yes |
| Single | Best | F6 | 1418 | *Anaerostipes hadrus* | 649756.24 | 98.5 | 93.9 | 100 | 12.1 | Yes |

| | | | | | | | | | | |
|---|---|---|---|---|---|---|---|---|---|---|
| Single | Best | F6 | 1637 | *Bacteroides caccae* | 47678.6 | 94.6 | 91.7 | 81.8 | 3.6 | Yes |
| Single | Best | F6 | 1152 | *Bacteroides thetaiotaomicron* | 818.287 | 97.6 | 91.8 | 90.7 | 15.1 | Yes |
| Single | Best | F6 | 1048 | *Faecalibacterium prausnitzii* | 853.27 | 98.3 | 93.9 | 98 | 19.1 | Yes |
| Paired | Best | F6 | 1391 | *Bacteroides cellulosilyticus* | 246787.5 | 97.7 | 93.7 | 91.5 | 10.9 | Yes |
| Paired | Best | F6 | 1764 | *Bacteroides fragilis* | 817.199 | 95.8 | 93.8 | 92.2 | 3.6 | Yes |
| Paired | Best | F6 | 2015 | *Odoribacter splanchnicus* | 28118.4 | 99.6 | 98.7 | 100 | 1.2 | Yes |
| Paired | Best | F6 | 2056 | *Parabacteroides distasonis* | 823.7 | 99.1 | 96.9 | 100 | 0 | Yes |
| Single | Best | F9 | 1903 | *Campylobacter hominis* | 360107.7 | 99.6 | 99.1 | 97.3 | 3 | Yes |
| Single | All | F9 | 1093 | *Proteus mirabilis* | 584.299 | 98.1 | 86.1 | 100 | 16.9 | Yes |
| Single | All | F9 | 1620 | *Pseudomonas aeruginosa* | 287.3918 | 98.9 | 93.9 | 99.7 | 8 | Yes |
| Paired | Best | G0 | 1871 | *Akkermansia muciniphila* | 239935.85 | 99.9 | 99.2 | 100 | 4.2 | Yes |
| Single | Best | G0 | 2022 | *Bacteroides fragilis* | 1339290.3 | 99.9 | 98.4 | 100 | 1 | Yes |
| Single | Best | G0 | 1950 | *Bacteroides vulgatus* | 821.84 | 99.7 | 99.2 | 99.4 | 2.5 | Yes |
| Single | Best | G0 | 1765 | *Bifidobacterium catenulatum* | 1686.6 | 99 | 97.3 | 100 | 5.9 | Yes |
| Single | Best | G0 | 2049 | *Escherichia coli* | 562.23525 | 99.6 | 98.1 | 100 | 0.4 | Yes |
| Single | Best | G0 | 1994 | *Methanobrevibacter smithii* | 911133.5 | 98.8 | 98.6 | 100 | 1.6 | Yes |
| Paired | Best | G3 | 1448 | *Finegoldia magna* | 1260.9 | 99.6 | 97.1 | 100 | 12.2 | Yes |
| Paired | Best | G3 | 1934 | *Prevotella denticola* | 28129.7 | 98.5 | 97.2 | 100 | 2.5 | Yes |
| Single | All | G3 | 1670 | *Akkermansia muciniphila* | 239935.85 | 100 | 97.3 | 100 | 7.8 | Yes |
| Single | Best | G3 | 1875 | *Campylobacter hominis* | 360107.7 | 99.3 | 98.4 | 97.3 | 3.4 | Yes |
| Single | Best | G3 | 1950 | *Lawsonella clevelandensis* | 1528099.5 | 95.9 | 95.7 | 90.7 | 0 | Yes |
| Single | Best | G3 | 1858 | *Mobiluncus curtisii* | 887899.3 | 95.9 | 94.8 | 100 | 3.5 | Yes |

| | | | | | | | | | | |
|---|---|---|---|---|---|---|---|---|---|---|
| Single | Best | G3 | 1765 | *Negativicoccus massiliensis* | 1702287.3 | 96.6 | 95 | 100 | 5.4 | Yes |
| Paired | Best | G9 | 1826 | *Akkermansia muciniphila* | 239935.85 | 100 | 97.8 | 100 | 4.8 | Yes |
| Paired | Best | G9 | 1945 | *Anaerococcus prevotii* | 879305.3 | 98.9 | 98.2 | 100 | 2.5 | Yes |
| Paired | Best | G9 | 2009 | *Campylobacter hominis* | 360107.7 | 99.7 | 99.2 | 97.3 | 0.9 | Yes |
| Paired | Best | G9 | 1365 | *Campylobacter ureolyticus* | 827.23 | 96.6 | 94 | 95.6 | 12.3 | Yes |
| Paired | Best | G9 | 1550 | *Corynebacterium jeikeium* | 38289.26 | 96.7 | 93.6 | 99.5 | 9.3 | Yes |
| Paired | Best | G9 | 1627 | *Corynebacterium urealyticum* | 43771.9 | 96.7 | 93.6 | 98.7 | 7.6 | Yes |
| Paired | Best | G9 | 1982 | *Finegoldia magna* | 1260.9 | 99.8 | 99.3 | 100 | 2 | Yes |
| Paired | Best | G9 | 1952 | *Lawsonella clevelandensis* | 1528099.5 | 96.1 | 95.9 | 90.7 | 0 | Yes |
| Paired | Best | G9 | 1952 | *Mobiluncus curtisii* | 887899.3 | 96.2 | 95.6 | 100 | 1.8 | Yes |
| Paired | Best | G9 | 942 | *Peptoniphilus harei* | 54005.7 | 95.8 | 91.4 | 93.1 | 19.7 | Yes |
| Paired | Best | G9 | 1800 | *Proteus mirabilis* | 584.9 | 98.5 | 94.1 | 100 | 4.5 | Yes |
| Paired | Best | H0 | 2009 | *Escherichia coli* | 562.28156 | 99.6 | 98.1 | 100 | 1.2 | Yes |
| Single | All | H0 | 1007 | *Fusobacterium nucleatum* | 469603.3 | 98.2 | 90.2 | 99.4 | 19.4 | Yes |
| Single | Best | H0 | 1423 | *Acidaminococcus intestini* | 1120921.3 | 97 | 93.7 | 98.7 | 11.7 | Yes |
| Single | Best | H0 | 1268 | *Aerococcus viridans* | 1377.13 | 96.3 | 93 | 93 | 13.5 | Yes |
| Single | Best | H0 | 1148 | *Anaerococcus prevotii* | 879305.3 | 98.8 | 93.4 | 100 | 17.4 | Yes |
| Single | Best | H0 | 1800 | *Bacteroides fragilis* | 1339290.3 | 98.4 | 95.9 | 100 | 4.9 | Yes |
| Single | Best | H0 | 1939 | *Bacteroides vulgatus* | 821.509 | 98.2 | 97 | 96.2 | 1.6 | Yes |
| Single | Best | H0 | 1886 | *Finegoldia magna* | 1260.1 | 99.6 | 98.3 | 100 | 3.7 | Yes |
| Single | Best | H0 | 2063 | *Proteus mirabilis* | 1125694.3 | 98.8 | 97.6 | 100 | 0 | Yes |
| Paired | All | H6 | 1777 | *Aerococcus viridans* | 1377.13 | 97.2 | 95 | 93.7 | 3.9 | Yes |
| Paired | All | H6 | 2068 | *Enterococcus faecalis* | 1158622.3 | 99.7 | 98 | 100 | 0 | Yes |
| Paired | Best | H6 | 1734 | *Escherichia coli* | 562.28156 | 99.3 | 93.1 | 100 | 5.6 | Yes |
| Single | Best | H6 | 1581 | *Lactobacillus crispatus* | 575597.3 | 96.2 | 94.3 | 89.9 | 6.9 | Yes |

| | | | | | | | | | | |
|---|---|---|---|---|---|---|---|---|---|---|
| Single | Best | H6 | 1566 | *Staphylococcus aureus* | 1280.10924 | 95.6 | 92.8 | 93 | 7.5 | Yes |
| Single | Best | H6 | 1759 | *Staphylococcus haemolyticus* | 1283.114 | 94.8 | 93.2 | 95.9 | 4.3 | Yes |
| Paired | Best | I0 | 1618 | *Finegoldia magna* | 1260.14 | 99.2 | 96.2 | 100 | 8.6 | Yes |
| Paired | Best | I0 | 1864 | *Peptoniphilus harei* | 54005.7 | 97.4 | 95.8 | 100 | 3.6 | Yes |
| Paired | Best | I0 | 1862 | *Proteus mirabilis* | 584.9 | 98.8 | 94.7 | 100 | 3.4 | Yes |
| Paired | Best | I3 | 1732 | *Odoribacter splanchnicus* | 28118.4 | 99.5 | 96.1 | 100 | 6.3 | Yes |
| Single | All | I3 | 1745 | *Methanobrevibacter smithii* | 2173.7 | 99.6 | 97.3 | 100 | 6.3 | Yes |
| Single | Best | I3 | 1486 | *Bacteroides caccae* | 47678.171 | 97.6 | 94.2 | 89 | 8.6 | Yes |
| Single | Best | I3 | 1945 | *Campylobacter hominis* | 360107.7 | 99.6 | 99.3 | 97.3 | 2.2 | Yes |
| Single | Best | I3 | 2008 | *Escherichia coli* | 562.30949 | 99.6 | 97.9 | 99.6 | 1.1 | Yes |
| Single | Best | I3 | 1554 | *Parabacteroides distasonis* | 823.236 | 99.4 | 94 | 100 | 9.4 | No |
| Single | Best | I6 | 1659 | *Collinsella aerofaciens* | 74426.49 | 97.1 | 92.9 | 99.2 | 6.9 | Yes |
| Single | Best | I6 | 2088 | *Methanobrevibacter smithii* | 2173.7 | 99.9 | 99.9 | 100 | 0 | Yes |
| Paired | Best | I9 | 1991 | *Campylobacter hominis* | 360107.7 | 99.6 | 99.4 | 97.3 | 1.3 | Yes |
| Single | Best | I9 | 1524 | *Bacteroides thetaiotaomicron* | 818.2 | 99.6 | 96.8 | 99.9 | 10.6 | Yes |
| Single | Best | I9 | 2021 | *Parabacteroides distasonis* | 823.236 | 99.2 | 97.8 | 100 | 0.9 | Yes |
| Paired | Best | I12 | 2081 | *Odoribacter splanchnicus* | 28118.4 | 99.9 | 99.2 | 100 | 0 | Yes |
| Paired | Best | I12 | 1862 | *Parabacteroides distasonis* | 823.236 | 99.3 | 95.6 | 100 | 3.6 | Yes |
| Paired | Best | I12 | 1891 | *Eggerthella lenta* | 84112.14 | 98.9 | 96.9 | 100 | 3.3 | Yes |
| Paired | Best | I12 | 2069 | *Escherichia coli* | 562.23525 | 99.6 | 98.1 | 100 | 0 | Yes |
| Single | All | I12 | 1188 | *Phascolarctobacterium faecium* | 1122957.3 | 94.9 | 88.2 | 98.7 | 15.2 | Yes |
| Single | Best | I12 | 1747 | *Bacteroides caccae* | 47678.171 | 98.9 | 97.2 | 95.8 | 5.4 | Yes |
| Single | Best | I12 | 2068 | *Bacteroides fragilis* | 1339290.3 | 99 | 98 | 100 | 0 | Yes |

| | | | | | | | | | | |
|---|---|---|---|---|---|---|---|---|---|---|
| Single | Best | I12 | 1880 | *Bacteroides thetaiotaomicron* | 818.2 | 99 | 97.5 | 96.3 | 2.9 | Yes |
| Single | Best | I12 | 867 | *Bifidobacterium bifidum* | 1681.45 | 95.6 | 90.8 | 85.8 | 19.6 | Yes |
| Single | Best | I12 | 2003 | *Candidatus Methanomassiliicoccus* | 1295009.4 | 97 | 96.2 | 95.5 | 0 | Yes |
| Single | Best | I12 | 2088 | *Methanobrevibacter smithii* | 2173.7 | 99.9 | 99.9 | 100 | 0 | Yes |
| Single | Best | J0 | 1929 | *Campylobacter ureolyticus* | 827.18 | 96.8 | 96.7 | 96.5 | 1.8 | Yes |
| Single | Best | J0 | 1233 | *Prevotella denticola* | 28129.7 | 98.4 | 95.3 | 100 | 16.1 | Yes |
| Single | Best | J0 | 1941 | *Proteus mirabilis* | 584.664 | 97.2 | 95.6 | 94.9 | 1 | Yes |
| Single | Best | J0 | 1460 | *Streptococcus oralis* | 1303.283 | 99.1 | 95.6 | 99.8 | 11.6 | Yes |
| Single | Best | J3 | 1187 | *Campylobacter hominis* | 360107.7 | 99.5 | 95.6 | 93 | 15.7 | Yes |
| Single | Best | J6 | 1912 | *Acidaminococcus intestini* | 1120921.3 | 97.4 | 96.3 | 99.3 | 2.6 | Yes |
| Single | Best | J6 | 1944 | *Campylobacter hominis* | 360107.7 | 99.6 | 99.2 | 97.3 | 2.2 | Yes |
| Single | Best | J6 | 1786 | *Escherichia coli* | 749531.3 | 99.6 | 94.2 | 100 | 4.8 | Yes |
| Paired | Best | J9 | 2007 | *Campylobacter hominis* | 360107.7 | 99.6 | 99 | 97.3 | 0.9 | Yes |
| Paired | Best | J9 | 1157 | *Corynebacterium urealyticum* | 43771.9 | 97 | 93.2 | 99.2 | 17 | Yes |
| Paired | Best | J12 | 1461 | *Parabacteroides distasonis* | 1339341.3 | 98.9 | 94.2 | 99 | 11.1 | Yes |
| Single | All | J12 | 1266 | *Campylobacter ureolyticus* | 827.18 | 95.7 | 92.6 | 92.2 | 13.3 | Yes |
| Single | All | J12 | 1286 | *Collinsella aerofaciens* | 74426.49 | 97.4 | 92.9 | 96.9 | 13.9 | Yes |
| Single | Best | J12 | 1882 | *Campylobacter hominis* | 360107.7 | 99.5 | 99 | 97.3 | 3.4 | Yes |
| Paired | Best | K0 | 1923 | *Campylobacter hominis* | 360107.7 | 99.6 | 99.1 | 97.3 | 2.6 | Yes |
| Single | Best | K0 | 1611 | *Peptoniphilus harei* | 54005.7 | 96.2 | 94.2 | 95.5 | 7.4 | Yes |
| Paired | Best | K3 | 1083 | *Bacteroides vulgatus* | 821.509 | 99.6 | 93.9 | 100 | 18.8 | Yes |

| | | | | | | | | | | |
|---|---|---|---|---|---|---|---|---|---|---|
| Paired | Best | K3 | 1891 | *Bifidobacterium bifidum* | 1681.55 | 99.3 | 98.1 | 99.7 | 3.5 | Yes |
| Paired | Best | K3 | 1723 | *Bifidobacterium longum* | 216816.147 | 99.4 | 97.9 | 98.6 | 6.6 | Yes |
| Single | Best | K3 | 2002 | *Candidatus Methanomassiliicoccus* | 1295009.4 | 97 | 96.1 | 95.5 | 0 | Yes |
| Single | Best | K3 | 2030 | *Escherichia coli* | 562.28156 | 99.9 | 98.1 | 99.6 | 0.7 | Yes |
| Single | Best | K3 | 1575 | *Eubacterium callanderi* | 53442.4 | 98.9 | 96.4 | 98 | 9.1 | Yes |
| Single | Best | K3 | 2088 | *Methanobrevibacter smithii* | 2173.71 | 99.9 | 99.9 | 100 | 0 | Yes |
| Single | Best | K6 | 1898 | *Bacteroides caccae* | 47678.175 | 98.7 | 97.4 | 96.7 | 2.6 | Yes |
| Single | Best | K6 | 1976 | *Odoribacter splanchnicus* | 28118.38 | 99.7 | 98.8 | 100 | 2 | Yes |
| Single | Best | K9 | 1725 | *Campylobacter hominis* | 360107.7 | 99.3 | 97.9 | 97.3 | 6.3 | Yes |
| Single | Best | K9 | 1375 | *Campylobacter ureolyticus* | 827.23 | 95.6 | 92.8 | 93.9 | 11.5 | Yes |

*Additional Table 4: Bins selected for DeepARG analysis*

This table lists all of the bins generated by PATRIC that were selected based on the criteria in Additional Table 3 to be analyzed using DeepARG. It includes all quality scores used to assess bin quality, as well as the PATRIC reference genome used to annotate the bin.

| Sample | SPREAD ID | BioSample ID | Sample | SPREAD ID | BioSample ID | Sample | SPREAD ID | BioSample ID |
|---|---|---|---|---|---|---|---|---|
| **1_A2** | 02/007/3/6/R | S02_007_3_6_R | **2_B4** | 09/018/B/6/R | S09_018_B_6_R | **3_B8** | 26/031/6/5/R | S26_031_6_5_R |
| **1_A8** | 02/021/6/8/R | S02_021_6_8_R | **2_B11** | 09/048/9/5/R | S09_048_9_5_R | **3_B11** | 26/038/3/9/R | S26_038_3_9_R |
| **1_A11** | 02/023/3/6/R | S02_023_3_6_R | **2_C1** | 09/085/3/5/R | S09_085_3_5_R | **3_C2** | 29/013/6/9/R | S29_013_6_9_R |
| **1_B3** | 02/032/B/7/R | S02_032_B_7_R | **2_C2** | 09/086/B/5/R | S09_086_B_5_R | **3_C4** | 31/039/B/7/R | S31_039_B_7_R |
| **1_B5** | 02/041/3/7/R | S02_041_3_7_R | **2_C5** | 09/099/9/9/R | S09_099_9_9_R | **3_C8** | 32/019/6/5/R | S32_019_6_5_R |
| **1_B11** | 04/003/6/5/R | S04_003_6_5_R | **2_C9** | 09/138/9/9/R | S09_138_9_9_R | **3_D2** | 32/022/12/5/R | S32_022_12_5_R |
| **1_C6** | 04/011/12/5/R | S04_011_12_5_R | **2_C12** | 09/143/9/9/R | S09_143_9_9_R | **3_D8** | 32/052/9/9/R | S32_052_9_9_R |
| **1_D2** | 04/059/6/9/R | S04_059_6_9_R | **2_D2** | 09/153/3/9/R | S09_153_3_9_R | **3_D10** | 34/009/B/2/R | S34_009_B_2_R |
| **1_D8** | 06/007/9/6/R | S06_007_9_6_R | **2_D6** | 09/187/3/9/R | S09_187_3_9_R | **3_E4** | 35/010/B/5/R | S35_010_B_5_R |
| **1_D12** | 06/027/6/6/R | S06_027_6_6_R | **2_D10** | 09/192/6/9/R | S09_192_6_9_R | **3_E6** | 35/031/B/9/R | S35_031_B_9_R |
| **1_E4** | 06/040/6/7/R | S06_040_6_7_R | **2_E1** | 09/214/6/9/R | S09_214_6_9_R | **3_E8** | 36/007/B/7/R | S36_007_B_7_R |
| **1_E9** | 06/048/9/5/R | S06_048_9_5_R | **2_E5** | 10/010/6/6/R | S10_010_6_6_R | **3_F4** | 38/001/3/5/R | S38_001_3_5_R |
| **1_F1** | 06/060/6/5/R | S06_060_6_5_R | **2_F1** | 10/012/6/6/R | S10_012_6_6_R | **3_F6** | 38/004/3/5/R | S38_004_3_5_R |
| **1_F8** | 06/068/6/5/R | S06_068_6_5_R | **2_F11** | 13/030/3/6/R | S13_030_3_6_R | **3_F10** | 38/017/B/5/R | S38_017_B_5_R |
| **1_F10** | 06/071/B/6/R | S06_071_B_6_R | **2_G6** | 13/035/12/7/R | S13_035_12_7_R | **3_G3** | 38/024/9/9/R | S38_024_9_9_R |
| **1_G3** | 06/083/9/9/R | S06_083_9_9_R | **2_G9** | 13/080/6/7/R | S13_080_6_7_R | **3_G7** | 39/008/6/7/R | S39_008_6_7_R |
| **1_G9** | 06/085/12/9/R | S06_085_12_9_R | **2_H2** | 19/009/6/5/R | S19_009_6_5_R | **3_G11** | 39/011/6/7/R | S39_011_6_7_R |
| **1_G10** | 06/102/B/9/R | S06_102_B_9_R | **2_H4** | 19/031/B/5/R | S19_031_B_5_R | **3_H6** | 40/038/9/9/R | S40_038_9_9_R |
| **1_H3** | 06/107/B/9/R | S06_107_B_9_R | **2_H10** | 21/012/12/7/R | S21_012_12_7_R | **4_A3** | 40/044/6/9/R | S40_044_6_9_R |
| **1_H6** | 06/108/3/9/R | S06_108_3_9_R | **3_A7** | 21/037/6/7/R | S21_037_6_7_R | **4_A12** | 42/002/9/7R/2 | S42_002_9_7R_2 |
| **2_A2** | 07/020/3/7/R | S07_020_3_7_R | **3_A12** | 21/060/12/7/R | S21_060_12_7_R | **4_B4** | 42/014/6/7/R | S42_014_6_7_R |
| **2_A6** | 07/056/3/7/R | S07_056_3_7_R | **3_B4** | 23/025/9/9/R | S23_025_9_9_R | **4_B7** | 42/015/3/7/R | S42_015_3_7_R |
| **2_B2** | 07/059/6/7/R | S07_059_6_7_R | | | | | | |

*Additional Table 5: BioProject Sample Identifiers for Test Dataset*

This table lists the sample names used in this study, the SPREAD IDs, and the BioProject PRJNA531921 sample names for the shotgun metagenomics sequencing files of the 67-sample dataset used to test the multiple linear regression developed from the levofloxacin dataset.
