## Additional File 1 for "Antimicrobial resistance gene prevalence in a population of patients with advanced dementia is related to specific pathobionts"

#Rowan-Nash, Araos, D’Agata, & Belenky
#All analysis performed by ADR

#All metagenomic processing was run using the Brown University OSCAR computing cluster

**#KNEADDATA**

**#kneaddata v0.6.1 was used to remove contaminating human sequences from metagenomic sequence files**
**#The kneaddata "Homo_sapiens_Bowtie2_v0.1" database was utilized**

**#The following batch script was submitted to run kneaddata in parallel on files where paired reads were concatenated into single files:**

```
samples=( [SAMPLE NAMES )
f=${samples[$SLURM_ARRAY_TASK_ID]}
module load kneaddata/0.6.1
kneaddata --input [LOCATION]/${f}.fastq --reference-db
[LOCATION]/Homo_sapiens_Bowtie2_v0.1 --output [LOCATION]/knead
```

**#These files were then either used directly as single-read library input for metagenomic assembly in PATRIC, input for the metagenomics functional assignment pipeline HUMANn2 and antibiotic resistance analysis program DeepARG-SS, or split into forward and reverse reads and processed through Kraken2 and Bracken for taxonomic assignment.**

**#The following batch script was submitted to run kneaddata in parallel on paired read files:**

```
samples=( [SAMPLE NAMES )
f=${samples[$SLURM_ARRAY_TASK_ID]}
module load kneaddata/0.6.1
kneaddata --input [LOCATION]/${f}._R1.fastq [LOCATION]/${f}._R1.fastq --
reference-db [LOCATION]/Homo_sapiens_Bowtie2_v0.1 --output [LOCATION]/knead
```

**#These files were then used as paired-read library input for metagenomic assembly in PATRIC**

**#Kraken2 & Bracken2, Diversity Analysis in R**

**#Kraken2 and Bracken were installed and run in a conda environment, and wget had to be installed manually:**

```
conda config --add channels defaults
conda config --add channels bioconda
conda config --add channels conda-forge
conda create -n kraken kraken2 bracken
conda install wget --name kraken
```

**#First a custom database of bacterial and archaeal genomes was created using the following script:**

```
source activate kraken
kraken2-build --download-taxonomy --db kraken2db
kraken2-build --download-library bacteria --db kraken2db
kraken2-build --download-library archaea --db kraken2db
kraken2-build --build --db kraken2db
```

**#Then the kneaddata-processed sequence files were run through Kraken 2, using the bacterial/archaeal database, using the following script**

```
samples=( [SAMPLE NAMES] )
f=${samples[$SLURM_ARRAY_TASK_ID]}
source activate kraken
kraken2 --db [LOCATION]/database [LOCATION]/${f}_kneaddata.fastq --use-names
--report ${f}_kraken2
```

**#The output of Kraken2 was then run against Bracken to quantify relative abundances**
**#Used Kraken2 database to create a Bracken database file using the following script:**
**#Note1: due to using conda rather than standard install, a small change had to be made to the bracken-build script, to define the correct locatin of the kmer2read_distr script**
**#Note2: specified threads (-t), kmer length (-k, 35 standard for kraken2), and read length (150bp)**

```
source activate kraken
bracken-build -d [LOCATION]/database -t 10 -k 35 -l 150
```

**#Then used bracken to reestimate kraken2 results using the following script: (-r for read length, -l for level, -t for threads)**

```
samples=( [SAMPLE NAMES] )
f=${samples[$SLURM_ARRAY_TASK_ID]}
source activate kraken
bracken -d [DATABASE LOCATION] -i [LOCATION]/${f}_kraken2 -o
${f}.bracken.phylum -r 150 -l P -t 10
bracken -d [DATABASE LOCATION] -i [LOCATION]/${f}_kraken2 -o
${f}.bracken.genus -r 150 -l G -t 10
bracken -d [DATABASE LOCATION] -i [LOCATION]/${f}_kraken2 -o
${f}.bracken.species -r 150 -l S -t 10
```

```
#combined bracken files into a single output
bracken_combine_outputs.py --files *phylum.bracken -o bracken_phylum_all
bracken_combine_outputs.py --files *genus.bracken -o bracken_genus_all
bracken_combine_outputs.py --files *species.bracken -o bracken_species_all
#Converted bracken report files into .biom file for diversity analysis in
phyloseq (R)
kraken-biom *kraken2_bracken -o bracken.biom --fmt json

#In R, loaded required packages
library(phyloseq)
library(vegan)

#Imported biom file into phyloseq object
biomfilename = "bracken.biom"
data <- import_biom(biomfilename, parseFunction=parse_taxonomy_default)

#Estimated and exported alpha diversity
data.alpha<-estimate_richness(data)
write.csv(data.alpha, file="data-alpha-diversity.csv")

#Estimated and exported beta diversity (Bray-Curtis)
braycurtis <- phyloseq::distance(levo, method = "bray")
BCmat <- as.matrix(braycurtis)
write.csv(BCmat, file = "levo-braycurtis.csv")

#Created and exported PCoA values
braycurtis.pcoa <- ordinate(physeq = levo, method = "PCoA", distance =
"bray")
braycurtis.pcoa.export <- as.data.frame(braycurtis.pcoa$vectors, row.names =
NULL, optional = FALSE, cut.names = FALSE, col.names =
names(braycurtis.pcoa$vectors), fix.empty.names = TRUE, stringsAsFactors =
default.stringsAsFactors())
write.csv(braycurtis.pcoa.export, file="levo-braycurtis-pcoa.csv")
```

**#HUMANn2**
**#Used knead-processed files to analyze genes/pathways/etc using Humann2**
**conda create -n humann2 humann2**
**#Manually downloaded and installed metaphlan2 db_v20 into humann2/bin/**
**#Changed config files to reflect true database locations**

```
source activate humann2
humann2_config --update database_folders nucleotide [DATABASE LOCATION]/humann2_v0.11.2_dbs/chocophlan
humann2_config --update database_folders protein [DATABASE LOCATION]/humann2_v0.11.2_dbs/uniref
```

**#Ran humann2 using following script:**

```
samples=( [SAMPLE NAMES )
f=${samples[$SLURM_ARRAY_TASK_ID]}
source activate humann2
humann2 --input [KNEAD FILES LOCATION]/${f}_kneaddata.fastq --output [OUTPUT LOCATION]
```

**#Combined humann2 outputs into single tsv files**

```
humann2_join_tables --input [LOCATION] --output humann2_genefamilies.tsv --file_name genefamilies_relab
humann2_join_tables --input [LOCATION] --output humann2_pathcoverage.tsv --file_name pathcoverage
humann2_join_tables --input [LOCATION] --output humann2_pathabundance.tsv --file_name pathabundance_relab
```

**#Split tables into unstratified files**

```
humann2_split_stratified_table --input humann2_genefamilies.tsv –output [LOCATION]
humann2_split_stratified_table --input humann2_pathcoverage.tsv –output [LOCATION]
humann2_split_stratified_table --input humann2_pathabundance.tsv –output [LOCATION]
```

**#Regrouped humann2 gene families table into KEGG orthologs and GO terms**

```
humann2_regroup_table --input [LOCATION]/humann2_genefamilies.tsv --group uniref90_ko –output humann2_kegg.tsv
humann2_regroup_table --input [LOCATION]/humann2_genefamilies.tsv --group uniref90_go –output humann2_go.tsv
```

**#Renormalized tables from RPKs to relative abundances**

```
humann2_renorm_table --input [LOCATION]/humann2_kegg.tsv --output [LOCATION]/humann2_kegg_relab.tsv --units relab
humann2_renorm_table --input [LOCATION]/humann2_go.tsv --output [LOCATION]/humann2_go_relab.tsv --units relab
humann2_renorm_table --input [LOCATION]/humann2_pathabundance.tsv --output [LOCATION]/humann2_metacyc_relab.tsv --units relab
```

**#DeepARG**

**#Used deeparg to look for abundances and types of antibiotic resistance genes in the knead-processed reads of each sample using the following script**

```
samples=( [SAMPLE NAMES]
f=${samples[$SLURM_ARRAY_TASK_ID]}
module load deeparg/Jan2019
module load gcc/5.4
cd [DEEPARG LOCATION]
python deepARG.py --align --type nucl --reads --input [KNEAD FILES LOCATION]/${f}_kneaddata.fastq --output [OUTPUT FILES LOCATION]/${f}_args.out
```

**Used deeparg to look for abundances and types of antibiotic resistance genes in bins from PATRIC output using the following script**

```
samples=( [BIN NAMES] )
f=${samples[$SLURM_ARRAY_TASK_ID]}
module load deeparg/Jan2019
module load gcc/5.4
cd /users/arowan/data/Shared/deeparg-ss
python deepARG.py --align --type nucl --genes --input [BIN FILES LOCATION]/${f}.fastq --output [OUTPUT FILES LOCATION]/${f}_args.out
```

**#QIIME2 v2019.1**

**#QIIME2 loaded into conda environment**
```
source activate qiime2-2019.1
```

**#Imported data (demultiplexed files from Illumina)**
**#Note: Requires demultiplexed files and manifest file (.csv with 3 columns: sample-id, absolute-filepath, direction)**
```
qiime tools import --type 'SampleData[PairedEndSequencesWithQuality]' --
input-path manifest.csv --output-path demux.qza --input-format
PairedEndFastqManifestPhred33
```

**#Ran DADA2 to denoise data using the following script:**
```
module load anaconda/3-4.4.0
source activate qiime2-2017.8
qiime dada2 denoise-paired \
  --i-demultiplexed-seqs paired-end-demux.qza \
  --p-trim-left-f 30 \
  --p-trim-left-r 30 \
  --p-trunc-len-f 200 \
  --p-trunc-len-r 160
  --o-table table.qza \
  --o-representative-sequences rep-seqs.qza \
  --o-denoising-stats stats-dada2.qza
```

---

**#Generated phylogenetic trees**
```
qiime phylogeny align-to-tree-mafft-fasttree --i-sequences rep-seqs.qza --o-
alignment aligned-rep-seqs.qza --o-masked-alignment masked-alignment-rep-
seqs.qza --o-tree unrooted-tree.qza --o-rooted-tree rooted-tree.qza
```

**#Classified taxonomy**
```
qiime feature-classifier classify-sklearn --i-classifier silva-132-V4-
classifier.qza --i-reads rep-seqs.qza --o-classification taxonomy.qza
qiime metadata tabulate --m-input-file taxonomy.qza --o-visualization
taxonomy.qzv
```
